# CCT2 expression marks a translation-activated state in sporadic ALS motor neurons and is associated with functional decline

**DOI:** 10.64898/2026.10.06.757033

**Authors:** Priyanka Bhandary, Yizhou Wang, Mohamed Omar, Jason H. Moore, Zhiping Paul Wang

## Abstract

Sporadic disease accounts for about 90% of amyotrophic lateral sclerosis (ALS) cases, and its biological heterogeneity is a major contributor to the failure of trials that enroll patients under a single diagnosis. Because sporadic ALS lacks a shared genetic anchor, identifying biological signals is critical for effective patient stratification. We analyzed induced pluripotent stem cell–derived motor neurons from 619 AnswerALS participants using transcriptomics, proteomics, and whole-genome sequencing and identified CCT2 as the first single-gene marker of a molecular state within sporadic ALS, providing a measurable basis for trial enrollment according to biology. We first anchored the analysis in known ALS genotypes. C9orf72 and SOD1 motor neurons did not converge on a common signature but retained genotype-specific differences that bulk postmortem tissue compresses. Scoring 391 sporadic patients along three transcriptomic axes (bioenergetic stress, cell cycle, and inflammation) resolved three states. One state was marked by CCT2, a TRiC chaperonin subunit, which distinguished it from the quiescent state with an area under the curve of 0.892. CCT2 was also elevated at the protein level and associated with the rate of ALSFRS-R decline, although this association weakened after adjustment for culture composition. Upstream regulator analysis placed this state downstream of MYC and mTORC1 activation. No germline test across 360 whole genomes explained these states. Validation using Target ALS data showed that the relationship between CCT2 and the cell cycle program was also present in postmortem cortex enriched for layer 5b projection neurons.

## Introduction

ALS is a paralytic neurodegenerative disease. Also known as Lou Gehrig’s disease, it is a neurological disorder which targets the motor system (1–4). This disease sees the progressive degeneration of upper motor neurons (UMN) in the motor cortex and the lower motor neurons (LMN) in the brainstem and the spinal cord (3,4). ALS can be broadly classified into two forms: familial ALS (fALS) and sporadic ALS (sALS). fALS occurs in individuals with a family history of the disease, often indicating an inherited genetic predisposition, whereas sALS occurs in individuals with no known family history of ALS (5,6). Certain common mutations in fALS have been studied in extensive detail such as C9orf72 (6–11), SOD1 (12–16), ATXN2 (17–21), etc. Most cases, about 90 to 95%, are sporadic, and a small proportion are familial (22).

Studying sporadic ALS has been a challenge. Even though the definition states that patients with sALS have no family history, it is more complex than that (23). Sporadic ALS itself carries a substantial genetic component: twin studies put its heritability at roughly 60 percent (24), a figure closer to that seen in other complex, polygenic diseases than to a condition with no inherited contribution at all. What remains unresolved is which kind of genetic architecture explains that heritability. Some studies point to rare variants whose individual effects vary in strength, others to common variants each conferring only modest risk, and others still to some blend of the two, though none of these models has yet been confirmed with strong, reproducible evidence (25). Genes and variants have been linked to ALS through a range of techniques, but the strength of that evidence varies considerably, and several reported associations have failed to replicate (26,27). Interpreting any individual variant is harder still, since consensus criteria for scoring pathogenicity do not exist, and because ALS risk itself is age-dependent with many implicated variants showing incomplete penetrance, the same variants that show up in fALS pedigrees often turn up in unaffected control individuals as well (23). Some ALS cases may have a multigenic component, entail epigenetic alteration, or be caused by DNA damage, environmental risk factors, or viral infections, according to the diversity in penetrance and the absence of a single linked gene mutation in 85% of sALS (28).

There is heterogeneity in ALS, at both the biological and clinical level (29–31). At the clinical level, there is heterogeneity encountered with respect to site of onset, progression rate, and involvement of upper and lower motor neurons. It also overlaps significantly in terms of clinical symptoms with frontotemporal dementia (FTD). At the genetic level, there have been advances in discovering genes involved in ALS (32,33). Some of them have been associated directly and many others are seen as risk factors or disease modifier genes (33). Part of the genetic complexity comes from the fact that many of the discovered genes confer susceptibility rather than cause disease. Many studies also show that ALS associated genes show incomplete penetrance which is further complicated by the interaction of the genetic variants and effect of environment on them (29,33). This heterogeneity has a large impact on clinical trials. This heterogeneity is widely responsible for diluting treatment effects in ALS clinical trials, where biologically mixed populations get enrolled under one diagnosis (29,33–35). Several studies have argued for and demonstrated the case for molecular stratification as a strategy for ALS clinical trials (28,33,36–40). Stratifying ALS patients according to these molecular subtypes is a more promising strategy for success in long term clinical ALS trials (28,33,36).

Multiple studies have taken different approaches to molecular subtyping of ALS (37,39–43). Tam et al. in 2019 were the first to use unbiased machine learning algorithms to subtype ALS and discovered three subtypes – ALS-Ox, ALS-TE and ALS-Glia (42). Eshima et al. in 2023 confirmed these subtypes using unsupervised clustering by non-negative matrix factorization (NMF) (40). They were able to validate their subtypes with data from two separate repositories. Marriott et al. (2023) identified three subtypes in postmortem motor cortex that converge on similar biological themes, oxidative stress and apoptosis, neuroinflammation, and a synaptic and neuropeptide signaling program, and further showed that signatures derived from motor cortex could be detected, with high confidence, when projected onto peripheral blood from independent cohorts (39). Most recently, O’Neill et al. (2025) expanded the NYGC cohort to over 700 cortical and 600 spinal cord samples and confirmed the original three subtypes at single-nucleus resolution, showing that ALS-Glia cortex specifically loses layer 5 corticospinal projection neurons while the other subtypes do not (41). That independently derived clustering methods, applied to independent cohorts, repeatedly converge on the same small set of biological themes, oxidative and metabolic stress and neuroinflammation, is itself evidence that these are real, recurring axes of ALS heterogeneity rather than artifacts of any single cohort or method, and motivates the three-axis structure we use throughout this study.

A separate line of work has looked to peripheral blood as a more accessible alternative for de novo subtype discovery, rather than projecting an existing brain-derived signature onto it. Grima et al. (2023) profiled whole blood RNA sequencing from 96 sporadic ALS patients and 48 controls and identified four patient subgroups with distinct gene expression signatures (43). However, these blood-based subtypes tracked most closely with patient age and circulating immune cell composition rather than with clinical measures of disease progression, and a diagnostic classifier trained on this cohort saw its accuracy fall from 78% to 63% when tested on an independent blood dataset, with only a small fraction of differentially expressed genes replicating across studies. This contrasts with Marriott et al.’s projection-based approach above and suggests that blood may be more useful for detecting an already-validated brain signature than for discovering subtypes from scratch. Together, these efforts illustrate two distinct limitations facing molecular subtyping in ALS: postmortem tissue captures only the end stage of disease in a mixed cell population, while peripheral blood, though accessible throughout the disease course and useful for detecting known signatures, may principally reflect systemic immune status rather than motor neuron-intrinsic biology when used for de novo discovery. Neither approach provides direct access to the cell type primarily affected by disease.

Induced pluripotent stem cell lines (iPSCs) are particularly valuable for studying ALS. iPSCs are reprogrammed from somatic cells and can then be differentiated into various cell types. For studying ALS, motor neurons or glial cells can be produced from iPSCs. iPSCs were first produced using four transcription factors, reprogramming them from mouse and then human fibroblasts (44,45). An important advantage of iPSCs is that they can be derived directly from a patient’s own cells, retaining the complete genetic background (46). This would include any known causal mutation and the broader polygenic architecture underlying sporadic disease, while remaining scalable to support transcriptomic, proteomic, and genomic profiling of the same cells at cohort scale (47). This helps address the two limitations described above. Unlike peripheral blood, iPSC-MNs are the motor neuron itself, not a peripheral proxy for it, offering a clean cell type-specific population without the glial and immune contamination that dilutes signal in bulk tissue (46,48). Unlike postmortem tissue, they can be generated and studied from patients while alive, rather than being limited to a single end-stage glimpse collected after death (47). ALS subtyping has been done in bulk postmortem cortex and spinal cord, or in blood. In bulk postmortem motor cortex the transcriptomes of C9orf72, SOD1 and sporadic patients converge to the point of being hard to tell apart, with pairwise Spearman correlations of 0.49 to 0.75. The same three comparisons in purified motor neurons give 0.03 to 0.33. Glial and immune signal dominates bulk tissue and obscures the differences between genotypes, and presumably also the differences between sporadic patients. iPSC lines come from living patients earlier in the disease, so what we measure is not the end-stage common pathway that autopsy tissue necessarily captures. Matched whole genome sequencing on the same patients also lets us ask something the earlier studies could not: whether a transcriptomic subtype is encoded in germline DNA. We find that it is not in this cohort, and we therefore treat the state as regulatory rather than inherited. We also handle the partition differently. Unbiased clustering yields a label but no test of whether that label survives the technical variation of the platform that produced it. Ours is partly composition-dependent, so rather than defend the three-way split we move the central claim to a single gene that survives full adjustment. One gene is also easier to carry to another tissue or platform than a multi-gene signature. In our study, we used iPSCs from patient and control samples collected across different sites by the AnswerALS consortium (49).

Here, we test this directly using transcriptomic, proteomic, and whole genome sequencing data from iPSC-derived motor neurons in the AnswerALS cohort: 391 sporadic ALS patients, C9orf72 and SOD1 carriers, and healthy controls (49). C9orf72 motor neurons share part of their transcriptional program with sporadic ALS, while SOD1 motor neurons share little with either, so known ALS genotypes do not reduce to one common signature. This motivates the pathway framework we use, built from the pathways that separate the two familial genotypes. From there we define three pre-specified axes, metabolic, cell cycle, and inflammatory, and use them to identify three molecular states within sporadic disease. A series of sensitivity analyses complicates that picture: differentiation quality and cell composition explain part of the discrete three-way split. So we move the central claim down to the gene level, where CCT2 alone distinguishes a metabolically and cell-cycle dysregulated patient subgroup and survives adjustment for culture composition. We lay out a mechanistic hypothesis connecting CCT2 to coordinated mTOR-MYC translational activation. The resulting molecular state turns out to be largely independent of germline genetic variation, and its clinical associations are modest and confined to rates of functional decline. Last, the CCT2-to-cell-cycle relationship holds up in an independent tissue, sequencing modality, and cohort. Together, these results describe a form of ALS heterogeneity visible in patient-derived motor neurons, with one relationship, CCT2 to the cell cycle program, also present in postmortem cortex, and with implications for how future trials stratify patients.

## Results

### Study cohort and overall approach

We studied induced pluripotent stem cell derived motor neurons (iPSC-MNs) from the AnswerALS consortium, comprising 644 RNA sequencing samples from 619 participants: 400 samples from 391 unique sporadic ALS patients, 46 from 43 C9orf72 repeat expansion carriers, 13 from 12 SOD1 mutation carriers, and 185 from 173 controls (Table 1, Fig. 1).

**Figure 1.**
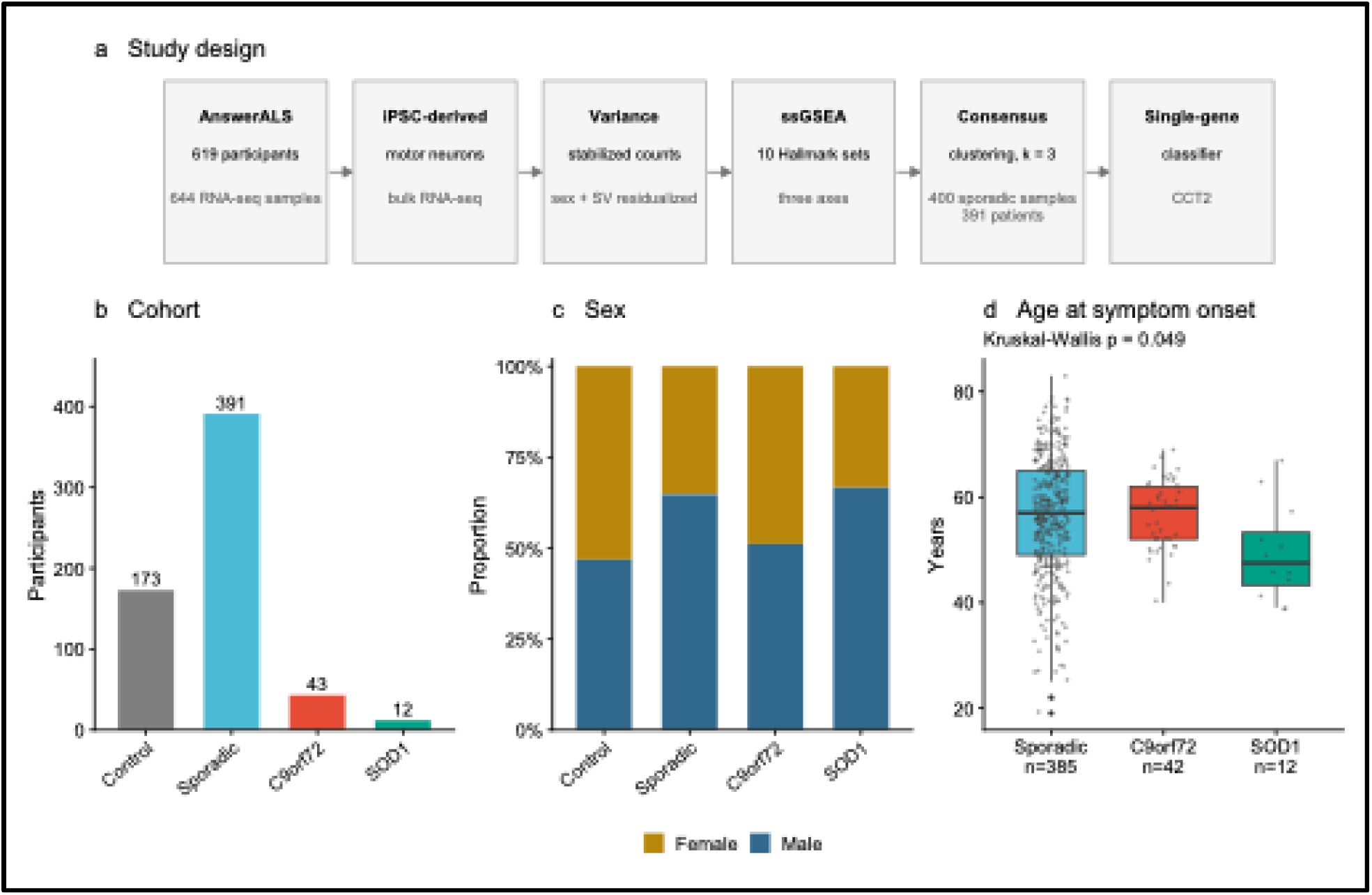
Study design and cohort. (a) Analysis workflow, from the AnswerALS cohort through variance-stabilized counts, ssGSEA scoring on ten Hallmark gene sets, consensus clustering of the 400 sporadic samples (391 patients), and single-gene classification. (b) Participants by group (173 control, 391 sporadic, 43 C9orf72, 12 SOD1; 644 RNA-seq samples in total). (c) Sex distribution by group. (d) Age at symptom onset, undefined for controls; boxes show median and interquartile range, points individual participants, Kruskal-Wallis P = 0.049. Numbers below each box are participants with a recorded onset age.

**Table 1.** Cohort characteristics.

| Characteristic | Control | Sporadic | C9orf72 | SOD1 |
| --- | --- | --- | --- | --- |
| Participants | 173 | 391 | 43 | 12 |
| RNA-seq samples | 185 | 400 | 46 | 13 |
| Female, n (%) | 92 (53) | 138 (35) | 21 (49) | 4 (33) |
| Male, n (%) | 81 (47) | 253 (65) | 22 (51) | 8 (67) |
| Age at symptom onset, years, mean (SD) | not applicable | 56.3 (11.2) | 56.8 (6.6) | 49.5 (9.1) |
| Age at first collection, years, mean (SD) | 55.1 (14.4) | 59.7 (10.8) | 59.0 (6.7) | 53.8 (7.7) |

Age at symptom onset is undefined for controls. Group assignment is from the AnswerALS mutation subtype field.

C9orf72 shares part of its transcriptional program with sporadic ALS, while SOD1 does not

Before focusing on sporadic disease, we asked how the two most common familial ALS genotypes in this cohort relate to sporadic ALS, since this comparison motivates treating sporadic ALS as biologically heterogeneous rather than as one uniform entity.

We fitted three differential expression contrasts against the same control group, one for each patient group, and compared the resulting log fold changes pairwise. Across all 24,253 expressed genes, the C9orf72 signature correlated with the sporadic signature at Spearman rho = 0.326 and with the SOD1 signature at 0.026, while SOD1 correlated with sporadic at 0.263. Genome-wide, the two familial genotypes are therefore similarly distant from sporadic disease; the difference between them appears when the comparison is anchored on the C9orf72 signature and at the pathway level. Anchoring on a single fixed reference set, the 500 genes most significantly changed in C9orf72, sharpened the contrast: those genes behaved concordantly in sporadic ALS (94.6% same direction, Spearman of fold changes = 0.803) but not in SOD1 (56.6%, rho = 0.295) (Fig. 2a-c; top-500 anchor in Methods)

**Figure 2.**
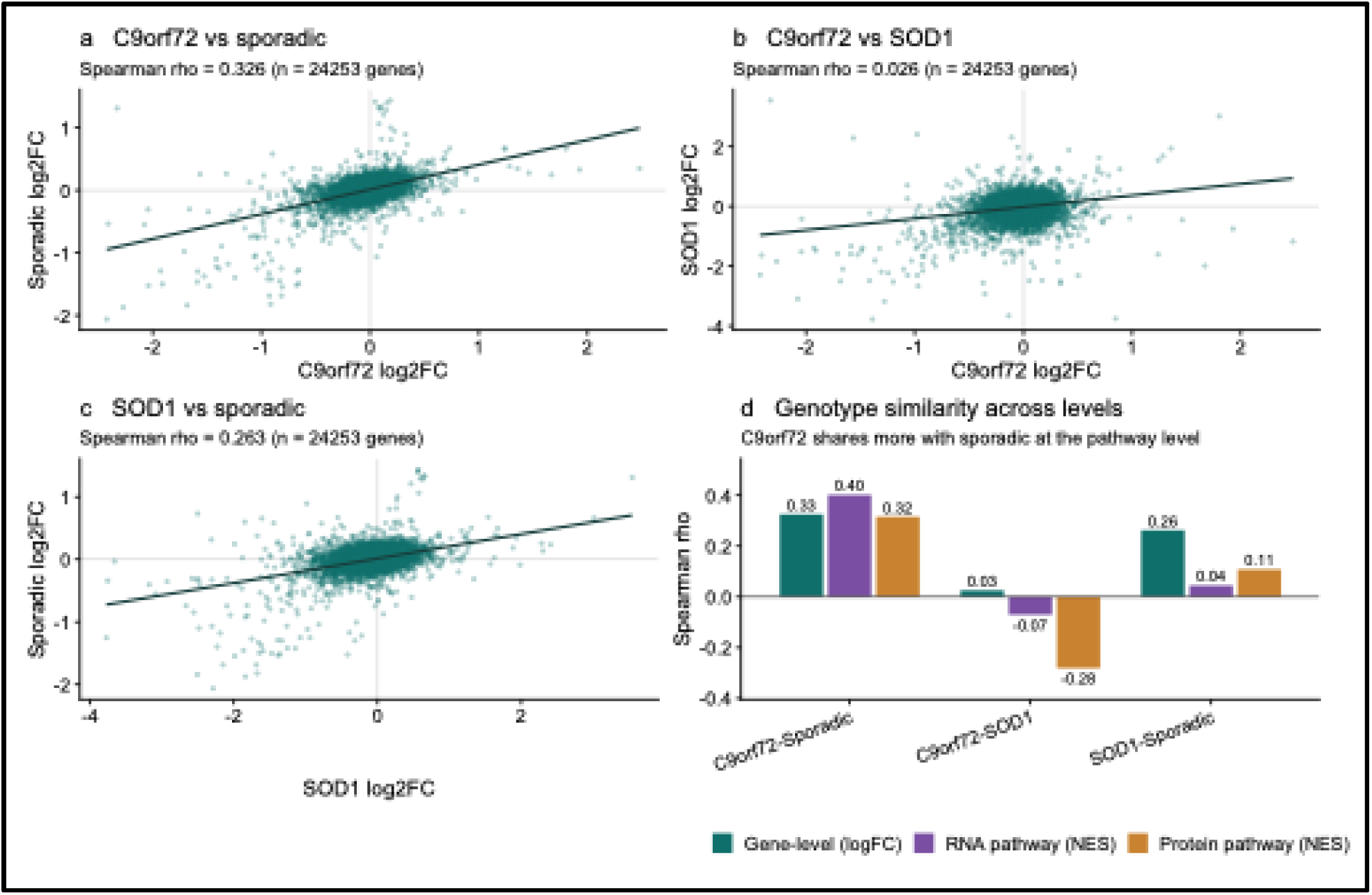
C9orf72 shares part of its transcriptional program with sporadic ALS, while SOD1 does not. (a-c) Gene-level differential expression against controls, compared pairwise across genotypes over all 24,253 expressed genes. Each point is one gene; the line is a linear fit and rho is the Spearman correlation of log2 fold changes. (d) The same three comparisons at three levels of aggregation: gene-level log2 fold changes across all expressed genes, and RNA and protein pathway normalized enrichment scores across all 50 Hallmark gene sets. RNA and protein rank the three comparisons in the same order.

At the pathway level the same ordering was seen. Hallmark normalized enrichment scores correlated at rho = 0.402 between C9orf72 and sporadic disease, at 0.045 between SOD1 and sporadic, and at −0.070 between C9orf72 and SOD1. This structure replicated independently in proteomics, using the identical construction across all 50 Hallmark pathways (C9orf72 versus sporadic rho = 0.319 with 66% sign agreement; SOD1 versus sporadic rho = 0.108, 58%; C9orf72 versus SOD1 rho = −0.282, 40%), arguing against an RNA-specific artifact (Fig. 2d).

Gene-level overlap between the RNA and protein layers could not be assessed, because no protein reached an adjusted P below 0.10 in any of the three contrasts (3,232 genes shared between the two platforms). All protein-level evidence presented in this paper is therefore pathway-level, where aggregate enrichment scores do not require individual proteins to reach significance. That is a depth limitation in this proteomics dataset, not a negative result about RNA-protein correspondence.

RNA and protein rank the three comparisons the same way. C9orf72 sits closest to sporadic disease, SOD1 close to neither, and C9orf72 against SOD1 falls at or below zero on both platforms. Familial and sporadic cases do not group by genotype. C9orf72 shares part of a transcriptional program with sporadic patients; SOD1 does something else. We take this as reason to treat sporadic ALS as a population with more than one molecular state in it, and it is why the axes defined below are built from pathways that separate these two familial genotypes.

As a control for whether this geometry simply reflects cell type composition, we computed the same contrasts in bulk postmortem motor cortex from the Target ALS cohort (43 control, 53 C9orf72, 18 SOD1, and 320 sporadic samples). There, all three cross-genotype gene-level correlations rose sharply (C9orf72 versus SOD1 rho = 0.487; C9orf72 versus sporadic rho = 0.751; SOD1 versus sporadic rho = 0.751), with directional agreement among the top 500 genes exceeding 97% in every comparison. Non-neuronal signal dominates bulk postmortem tissue and swamps the genotype differences a purified motor neuron preparation can still see. That is the argument for doing subtype discovery in iPSC-MNs.

### A pre-specified three-axis transcriptomic framework captures recurring pathway biology

Gene set enrichment across all three contrasts repeatedly nominated the same three biological themes: bioenergetic stress (oxidative phosphorylation, fatty acid metabolism, glycolysis), cell cycle and growth signaling (E2F targets, G2M checkpoint, MYC targets, DNA repair), and inflammation and injury (TNF alpha signaling via NF-kB, interferon gamma response, complement).

Motivated by this recurring structure, and by precedent from postmortem ALS subtyping (42), we defined three axes as unions of the relevant MSigDB Hallmark gene sets, fixed before any clustering was performed: AXIS_METABOLIC (512 genes), AXIS_CELL_CYCLE (610 genes), and AXIS_INFLAMMATION (538 genes)(50). Each patient received one single sample gene set enrichment (ssGSEA) score per axis (Fig. 3a). Axis scores did not differ across genotype groups (ANOVA P = 0.20, 0.09, and 0.53 for the metabolic, inflammation, and cell cycle axes respectively), so the axes describe variation within the cohort rather than separating familial from sporadic disease.

**Figure 3.**
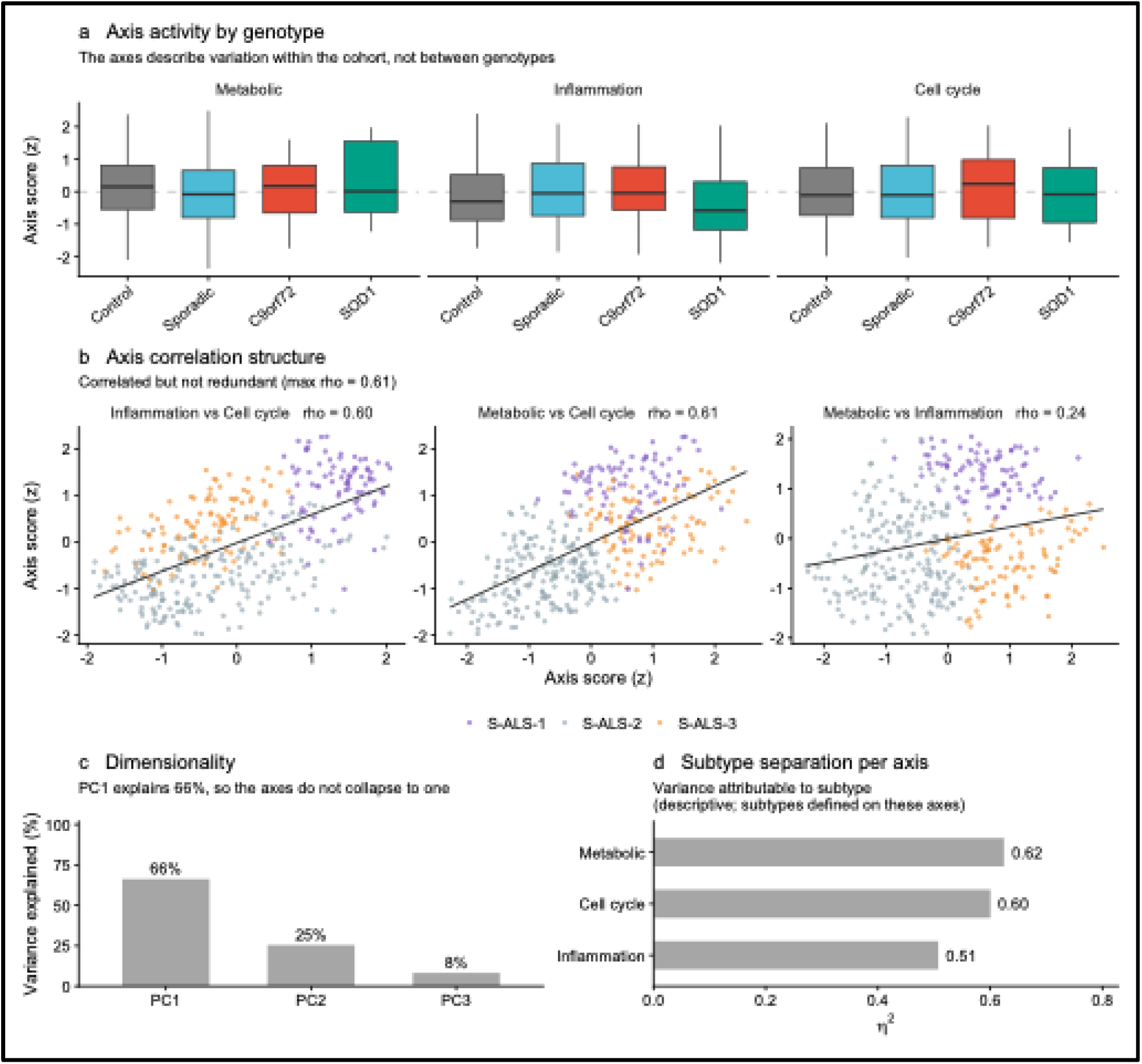
The three axes are correlated but separable. (a) Axis scores by genotype group. Scores do not differ across groups (ANOVA P = 0.20, 0.09 and 0.53 for the metabolic, inflammation and cell cycle axes), so the axes describe variation within the cohort rather than between genotypes. (b) Pairwise axis correlations across the 391 sporadic patients, colored by subtype. (c) Principal component analysis of the three axis scores. PC1 explains 66% of variance and loads positively on all three axes; PC2 explains 25% and contrasts the metabolic against the inflammation axis. PC1 loadings are 0.53, 0.53 and 0.66 for the metabolic, inflammation and cell cycle axes. (d) Variance in each axis attributable to subtype membership. Because subtypes were defined by clustering on these axes, these values describe how the partition distributes across the three dimensions and are not independent evidence that each axis contributes separately.

The three axes are correlated but not redundant. Pairwise Spearman correlations across the 391 sporadic patients were 0.24 between the metabolic and inflammation axes, 0.61 between metabolic and cell cycle, and 0.60 between inflammation and cell cycle (Fig. 3b). Because two of the three pairs exceed 0.5, we characterized the dimensionality directly rather than asserting independence. Principal component analysis of the three scores gave a first component explaining 66.3% of variance and loading positively on all three axes (0.53, 0.53, 0.66), which represents overall activation, and a second component explaining 25.4% and contrasting the metabolic against the inflammation axis with cell cycle at zero loading (Fig. 3c). Two components therefore capture 92% of the variance. The axes are separable but not independent: most of the variance is shared activation, with a metabolic against inflammatory contrast underneath it.

The three-axis structure was not an artifact of the Hallmark database: axes rebuilt from GO Biological Process and Reactome correlated with the Hallmark scores at r = 0.79 to 0.97 and produced partitions agreeing at adjusted Rand index 0.494 and 0.594, with balanced cluster sizes. A fully data-driven alternative based on thresholded gene lists recovered no significant GO term in any contrast (Supplementary Note 1).

### Consensus clustering identifies three transcriptomic states within sporadic ALS

Restricting to the 400 sporadic samples, we applied ConsensusClusterPlus to the three axis scores across k = 2 to 5, selecting k = 3 by minimizing the proportion of ambiguous clustering (PAC = 0.299 at k = 3, compared with 0.310 at k = 2, 0.368 at k = 4, and 0.356 at k = 5). We note that this minimum is shallow and that k = 2 is only marginally worse, so k = 3 is the best supported solution rather than a decisively separated one (Fig. 4a-b).

**Figure 4.**
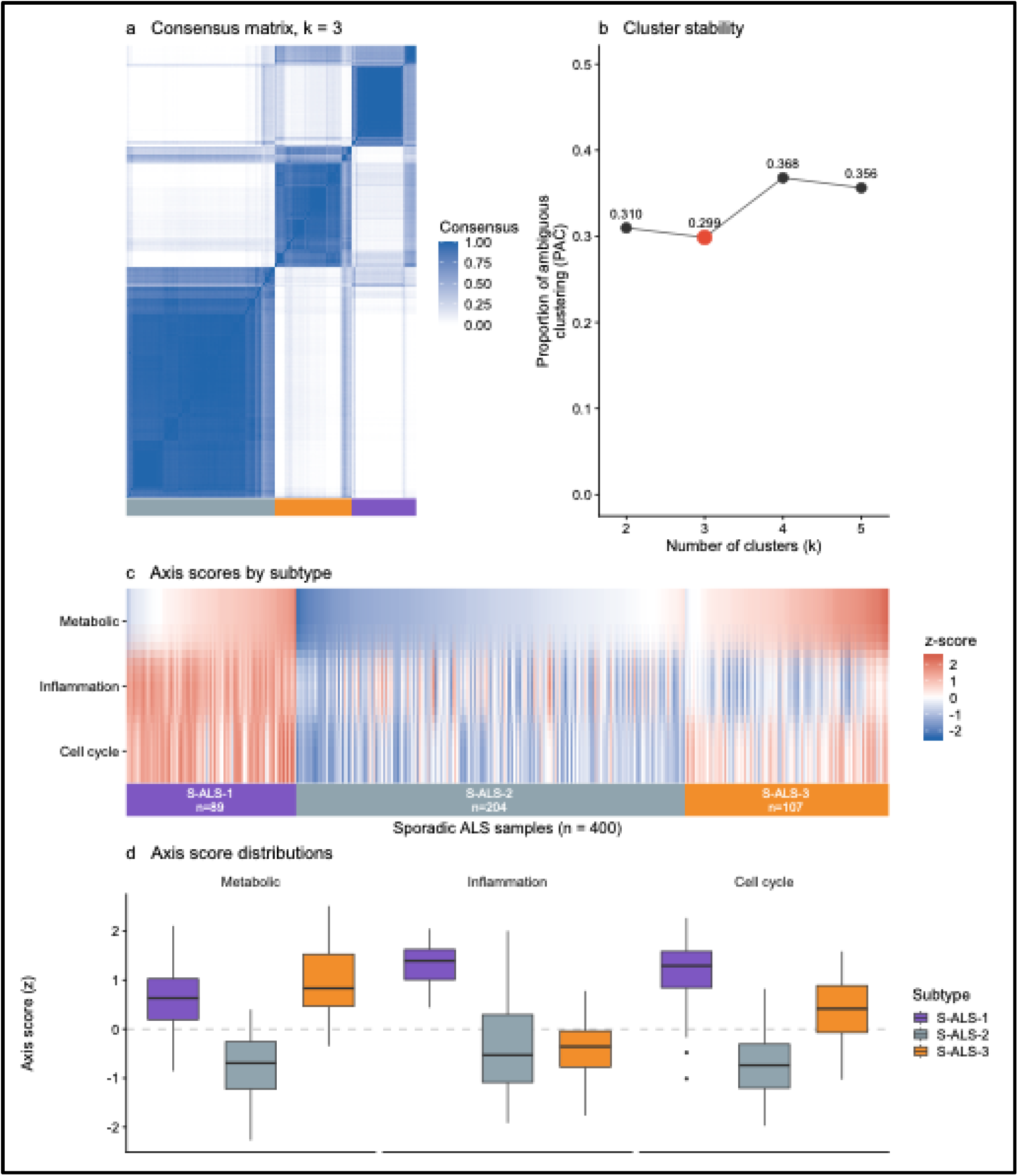
Consensus clustering of sporadic ALS. (a) Consensus matrix at k = 3 for the 400 sporadic samples, ordered by the consensus tree; the ribbon below marks subtype assignment. Clustering was performed at the sample level; the 391 patient-level labels used in all downstream analyses were taken from the earliest available sample per participant. (b) Proportion of ambiguous clustering across k. The minimum is at k = 3 (PAC = 0.299), only marginally below k = 2 (0.310), so k = 3 is the best supported solution rather than a decisively separated one. (c) Axis scores per sample, ordered by subtype and metabolic score; values were clipped at z = ±2.5. (d) Axis score distributions by subtype. S-ALS-1 is elevated on all three axes, S-ALS-2 is low on all three, and S-ALS-3 is elevated on the metabolic and cell cycle axes but below average on inflammation.

This partitioned sporadic patients into three states (Fig. 4c-d, Table 2): S-ALS-1 (89 samples, 85 patients, 22%), a pan-activated state elevated across all three axes and most strongly on inflammation (mean z = +0.60 metabolic, +1.34 inflammation, +1.17 cell cycle); S-ALS-2 (204 samples and patients, 52%), a quiescent state low across all three axes (−0.76, −0.37, −0.71), used as the reference group throughout; and S-ALS-3 (107 samples, 102 patients, 26%), elevated on the metabolic and cell cycle axes but below average on inflammation (+0.95, −0.41, +0.39).

**Table 2.** Characteristics of the three sporadic subtypes.

| Characteristic | S-ALS-1 | S-ALS-2 | S-ALS-3 |
| --- | --- | --- | --- |
| Patients, n (%) | 85 (22) | 204 (52) | 102 (26) |
| Metabolic axis, mean (SD) | 0.6 (0.6) | -0.8 (0.6) | 0.9 (0.6) |
| Inflammation axis, mean (SD) | 1.3 (0.4) | -0.4 (0.9) | -0.4 (0.6) |
| Cell cycle axis, mean (SD) | 1.2 (0.7) | -0.7 (0.6) | 0.4 (0.6) |
| CCT2, VST, mean (SD) | 13.01 (0.21) | 12.96 (0.15) | 13.20 (0.14) |
| Age at onset, years, mean (SD) | 56.6 (13.5) | 56.0 (11.0) | 56.6 (9.7) |
| Female, n (%) | 28 (33) | 65 (32) | 45 (44) |
| log10 NfL, mean (SD) | 1.6 (0.3) | 1.6 (0.3) | 1.7 (0.3) |
| ALSFRS-R baseline, mean (SD) | 33.1 (8.3) | 34.0 (8.9) | 34.0 (8.6) |
| ALSFRS-R slope, points/month, mean (SD) | -0.5 (0.6) | -0.6 (0.6) | -0.9 (0.8) |
| Differentiation quality, mean (SD) | -1.2 (1.4) | 0.2 (0.7) | 0.6 (0.5) |

Axis scores are z-scored across the 391 sporadic patients. Differentiation quality is the maturity marker composite minus the progenitor marker composite. Subtypes did not differ significantly by sex (chi-square = 4.74, df = 2, P = 0.094) or by age at onset. Differentiation quality differed across subtypes (eta squared = 0.382).

### Sensitivity analyses reposition the discrete three-way partition and validate the gene-level signal

Because iPSC differentiation efficiency varies across patient-derived lines, we ran three sensitivity checks of increasing stringency (Supplementary Note 2). Each repositioned the discrete partition: residualizing on a single glial marker gave an adjusted Rand index of 0.565 against the original labels, and residualizing on a full glial and differentiation-quality panel gave 0.273. Differentiation quality differed sharply across subtypes and the effect was dominated by S-ALS-1 (eta squared = 0.382), which we therefore treat as differentiation-quality confounded and demote to an exploratory category. S-ALS-3 versus S-ALS-2 is the primary comparison throughout and the less confounded one, although differentiation quality also differs modestly between these two subtypes (Table 2). The discrete label is substantially composition-dependent and cannot carry the paper on its own, so we use the three-axis framework as a discovery tool and put the primary claim at the gene level.

### CCT2 is a reliable single-gene marker of the S-ALS-3 state

CCT2, encoding the beta subunit of the TRiC/CCT chaperonin complex and annotated to the cell cycle axis through the HALLMARK_MYC_TARGETS_V1 gene set, separated S-ALS-3 from S-ALS-2 sharply (t = 14.0, P = 2.7 × 10^−32^; Wilcoxon P = 5.6 × 10^−29^), with a patient-level area under the ROC curve of 0.892 (bootstrap 95% confidence interval 0.851 to 0.927, 102 S-ALS-3 versus 204 S-ALS-2 patients) (Fig. 5a-c).

**Figure 5.**
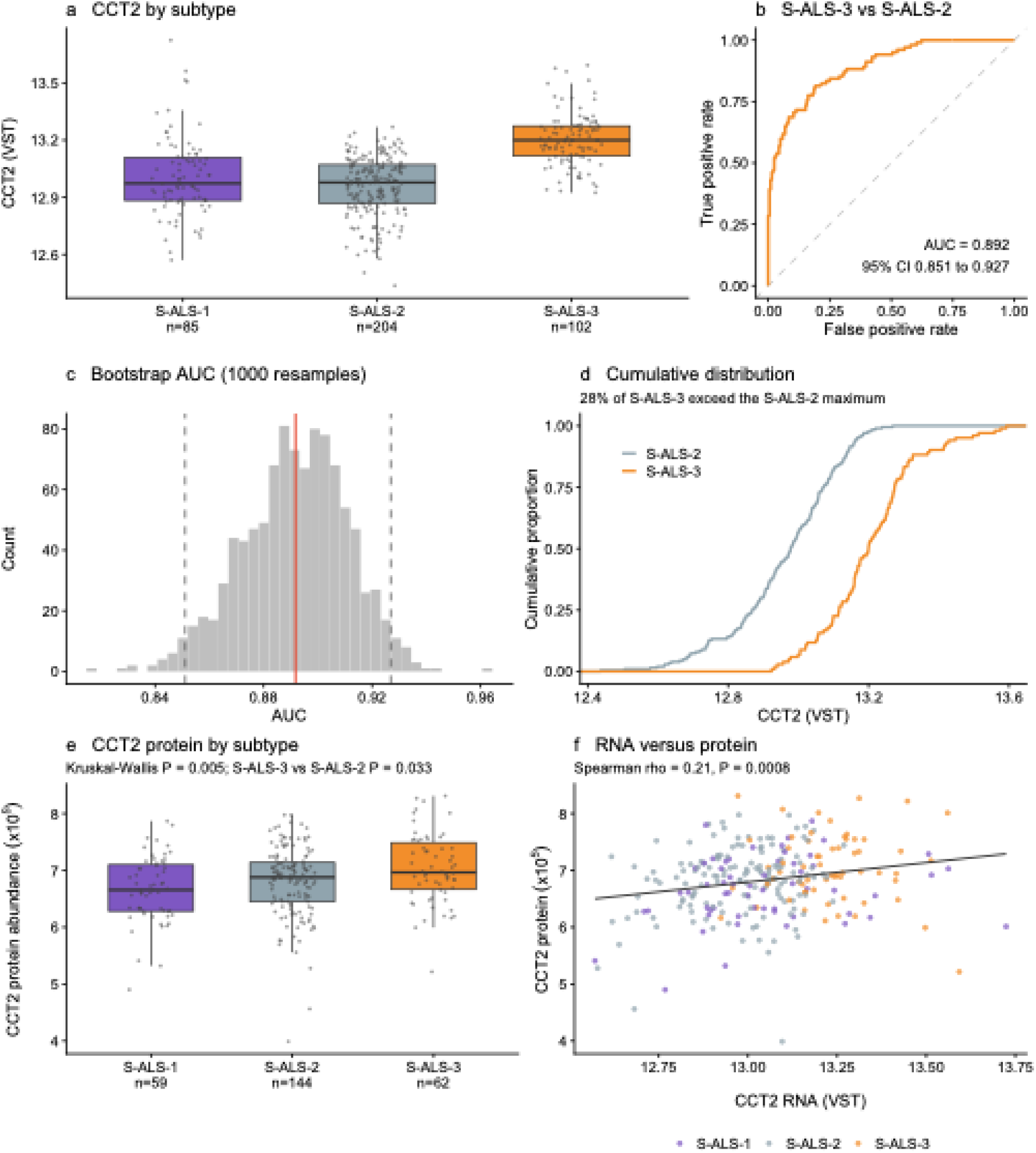
CCT2 marks the S-ALS-3 state at the RNA and protein level. (a) CCT2 expression by subtype in the 391 sporadic patients, one value per patient from the earliest available sample. (b) Receiver operating characteristic curve for CCT2 alone discriminating S-ALS-3 from S-ALS-2 (area under the curve 0.892, bootstrap 95% confidence interval 0.851 to 0.927). (c) Distribution of the area under the curve across 1,000 stratified bootstrap resamples; the red line marks the observed value and dashed lines the confidence interval. (d) Empirical cumulative distributions for the two subtypes. The separation is a uniform shift rather than an outlier effect: 28.4% of S-ALS-3 patients exceed the maximum S-ALS-2 value, Hartigan’s dip test finds no bimodality within S-ALS-3 (P = 0.679), and the area under the curve falls only to 0.873 when the 15 highest-CCT2 patients are removed. (e) CCT2 protein abundance by subtype in the 265 subtyped patients with proteomics. (f) CCT2 transcript expression versus protein abundance across the same patients.

This difference survived the culture-composition adjustments that repositioned the discrete partition. In a linear model, the unadjusted S-ALS-3 effect on CCT2 was +0.244 log2 units (P = 5.9 × 10^−33^). Adjusting for the proliferation, maturity, and differentiation quality composites reduced it only slightly (+0.217, P = 1.7 × 10^−24^), and further adding the full astrocyte, microglia, and oligodendrocyte panel did not attenuate it further (+0.231, P = 2.6 × 10^−28^). The gene-level signal therefore loses roughly a tenth of its magnitude under adjustments that substantially reshape the cluster assignment itself.

CCT2 elevation was a graded property of the S-ALS-3 population rather than an outlier effect. The distributions overlap substantially, with 28.4% of S-ALS-3 patients exceeding the maximum CCT2 value observed in S-ALS-2, and the separation reflects a uniform shift rather than a bimodal split: Hartigan’s dip test found no evidence of bimodality within S-ALS-3 (P = 0.679), and classification performance was barely affected by removing the 15 highest-CCT2 patients (area under the curve 0.873, compared with 0.892 in the full group) (Fig. 5d).

We then asked how minimal a gene panel could recover the subtype signal. One-vs-rest LASSO classifiers trained on the Hallmark axis genes recovered each subtype with compact panels of 12 to 16 genes (nested AUC 0.936 to 0.970; Supplementary Table S4). CCT2 was selected in 74.7% of bootstrap resamples for the S-ALS-3 classifier and was never selected for either other subtype, so it is specific to this state rather than a general marker of subtype membership.

Because the subtypes were defined within the sporadic cohort, controls took no part in the clustering and provide an external reference for CCT2. Relative to controls, CCT2 was lower in S-ALS-1 and S-ALS-2 and higher in S-ALS-3 (S-ALS-1 0.060 lower, P = 0.022; S-ALS-2 0.106 lower, P = 7.0 × 10^−10^; S-ALS-3 0.138 higher, P = 3.6 × 10^−12^; VST scale). The two directions cancel, so sporadic ALS as a whole differed from controls only marginally on this gene (0.033 lower, P = 0.044) while the difference between S-ALS-2 and S-ALS-3 within the same cohort was 0.244 (t = 14.0). A gene with no cohort-level signal therefore carries a strong signal once patients are stratified.

CCT2 elevation in S-ALS-3 replicated at the protein level. Among the 265 subtyped patients with proteomics (59 S-ALS-1, 144 S-ALS-2, 62 S-ALS-3), CCT2 protein differed across subtypes (Kruskal-Wallis P = 0.005) and was higher in S-ALS-3 than S-ALS-2 (Wilcoxon P = 0.033), and CCT2 transcript and protein were positively correlated across patients (Spearman rho = 0.21, P = 8 × 10^−4^). A composite score across all eight TRiC subunits did not differ between subtypes (P = 0.13), so the signal is specific to CCT2 and not a general change in chaperonin abundance (Fig. 5e-f).

### Upstream regulator analysis nominates a coordinated translation-activation mechanism

To generate a mechanistic hypothesis for the S-ALS-3 state, we performed upstream regulator and canonical pathway analysis (Ingenuity Pathway Analysis) on the S-ALS-3 versus S-ALS-2 differentially expressed gene list. The export returned 3,882 upstream regulators and 1,187 canonical pathways.

The regulator analysis nominated MYC as activated (activation z = +2.14, P = 6.0 × 10^−28^) and MYCN more strongly so (z = +5.58, P = 2.2 × 10^−19^). Two lines of evidence within the regulator analysis pointed to mTORC1 activation: the mTORC1 component RPTOR was predicted activated (z = +2.88, P = 1.4 × 10^−5^), while the mTOR inhibitors torin1 and sirolimus were predicted inhibited (z = −3.06, P = 2.5 × 10^−28^ and z = −3.93, P = 1.3 × 10^−21^), the expected signature of an activated pathway read through its pharmacological antagonists. LARP1 was predicted inhibited (z = −4.82, P = 1.9 × 10^−23^), consistent with release of 5’ TOP mRNA translation, a canonical LARP1-repressed program. RICTOR, the defining component of mTORC2, was predicted inhibited (z = −4.70, P = 3.1 × 10^−9^), indicating the signal is specific to mTORC1 rather than reflecting global mTOR activity (Fig. 6a, 6c). The regulators with the smallest P values overall were YAP1 and TGFB1, both predicted inhibited, together with several inflammatory regulators (IFNG, TNF, IL1B) also predicted inhibited, consistent with S-ALS-3 sitting below average on the inflammation axis (Fig. 6a).

**Figure 6.**
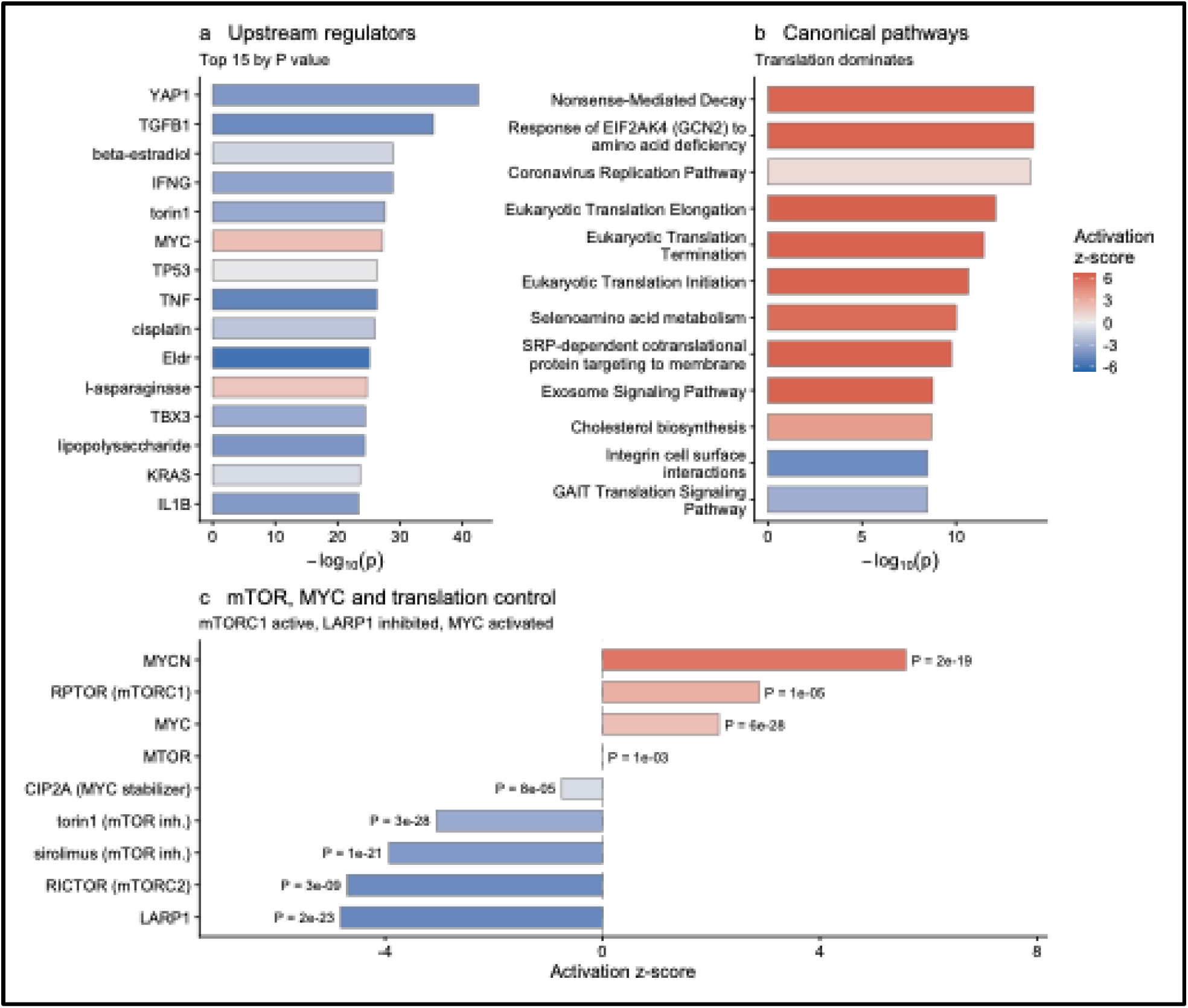
Upstream regulator and pathway analysis of S-ALS-3 versus S-ALS-2. (a) The 15 upstream regulators with the smallest P values, colored by predicted activation z-score. (b) The 12 canonical pathways with the smallest P values. Translation-related pathways dominate, together with the GCN2 (EIF2AK4) amino acid stress response and nonsense-mediated decay. (c) The mTOR, MYC and translation regulators specifically, ordered by activation z-score. MYC and MYCN are predicted activated and the mTORC1 component RPTOR is predicted activated, while the mTOR inhibitors torin1 and sirolimus are predicted inhibited, the expected signature of an active pathway read through its pharmacological antagonists. LARP1 inhibition is consistent with release of 5’ TOP mRNA translation. RICTOR inhibition indicates the signal is specific to mTORC1 rather than global mTOR activity. CIP2A is not predicted activated.

The top canonical pathways in the same contrast were dominated by translation, with eukaryotic translation elongation and termination among the most strongly activated (z = +6.56), alongside the GCN2 (EIF2AK4) amino acid deficiency response (z = +6.40, P = 7.3 × 10⁻¹) and nonsense-mediated decay (Fig. 6b). GO Biological Process over-representation analysis of the same gene list, run separately on genes higher and lower in S-ALS-3, recovered cytoplasmic translation as the top term among genes higher in S-ALS-3 (54 genes, adjusted P = 2.8 × 10^−21^), followed by cholesterol and sterol biosynthesis, aerobic respiration and canonical glycolysis. Genes lower in S-ALS-3 were enriched for miRNA-mediated gene silencing, mitotic division and spindle checkpoint, and cilium assembly terms (top adjusted P = 3.8 × 10^−19^). Scoring the four Hallmark sets that make up the cell cycle axis separately showed where the S-ALS-3 elevation comes from: MYC targets were higher in S-ALS-3 than in S-ALS-2 (Wilcoxon P = 2.1 × 10^−11^) and DNA repair modestly so (P = 0.002), E2F targets did not differ (P = 0.90), and G2M checkpoint genes were lower (P = 3.6 × 10^−4^). The cell cycle axis signal in S-ALS-3 is therefore a MYC target signal rather than a mitotic one, consistent with the regulator analysis and with the higher differentiation quality of S-ALS-3 cultures (Table 2).

The picture is a coordinated translation-activation program in S-ALS-3: mTORC1 activation with mTORC2 inhibition, MYC and MYCN activation driving transcription of ribosomal and translation machinery, LARP1 inhibition permitting translation of 5’ TOP transcripts, and GCN2-mediated amino acid stress signaling downstream of the resulting anabolic demand. We present this as a hypothesis supported by two enrichment methods applied to the same gene list, not as an experimentally validated mechanism. It would also explain why S-ALS-3 is elevated on both the metabolic and the cell cycle axis: both sit downstream of the same mTOR to MYC coordination.

### The subtype architecture is independent of germline variation

Given the strength of the transcriptomic signal, we asked whether subtype membership is encoded in germline DNA, using whole genome sequencing available for 360 sporadic patients. No test survived correction for multiple testing (Fig. 7, Table 3, Supplementary Table S3).

**Figure 7.**
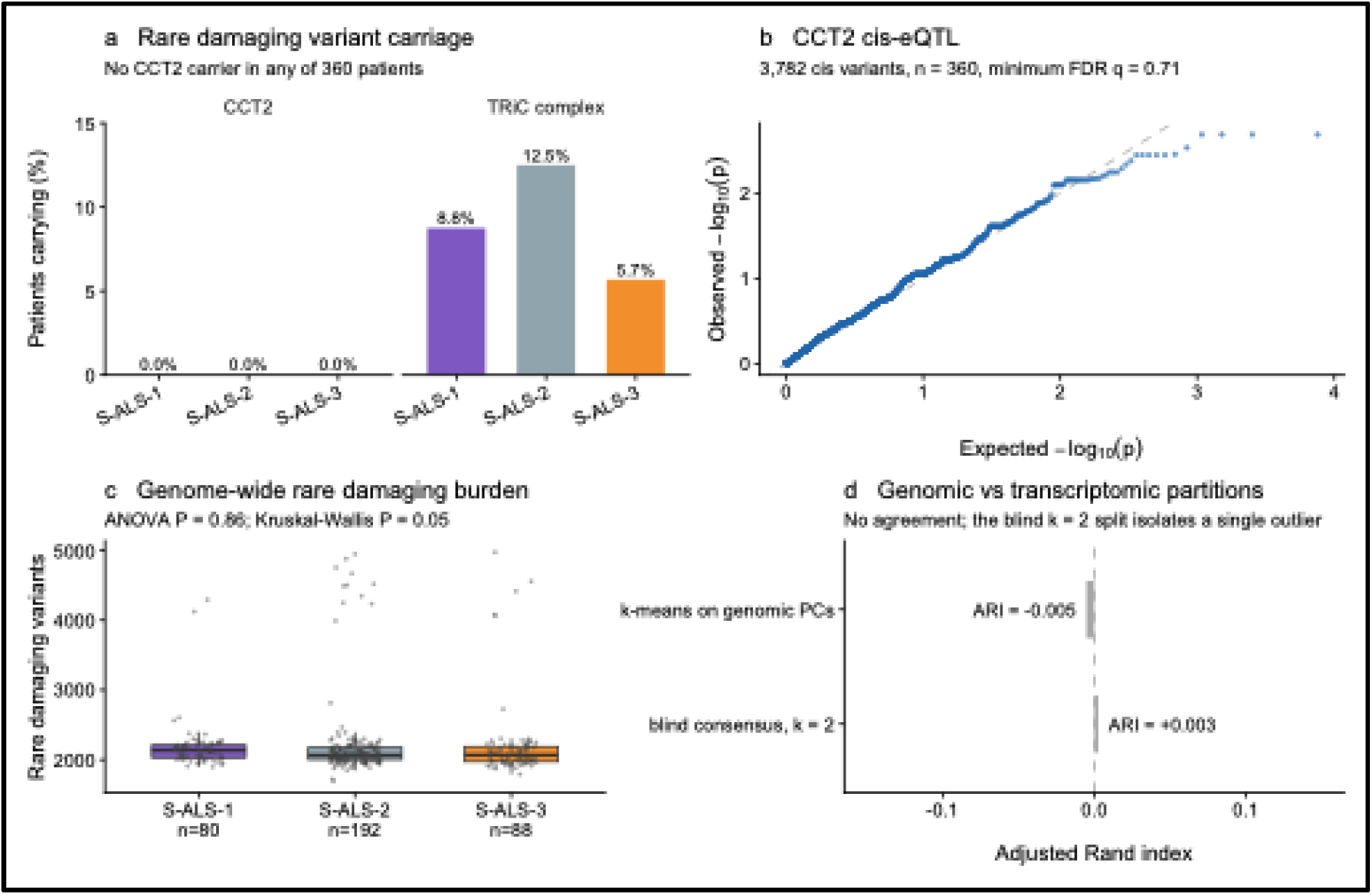
No evidence that the S-ALS-3/CCT2 state is explained by the germline variants tested. (a) Rare damaging variant carriage by subtype for CCT2 and for the eight-subunit TRiC complex. No CCT2 carrier was observed in any of the 360 sporadic patients with whole genome sequencing. (b) Quantile-quantile plot for the CCT2 cis-eQTL analysis across variants within 1 Mb of CCT2. (c) Genome-wide rare damaging variant burden by subtype. (d) Agreement between genomic and transcriptomic partitions. Neither k-means on genomic principal components nor blind consensus clustering recovers the transcriptomic labels; the blind k = 2 split separates a single high-burden outlier from the remaining 359 patients.

**Table 3.** Summary of germline genomic tests in 360 sporadic patients.

| Analysis | Test | Result |
| --- | --- | --- |
| Genome-wide rare damaging burden | ANOVA | P = 0.864 |
| Genome-wide rare damaging burden | Kruskal-Wallis | P = 0.05 |
| Genome-wide rare loss-of-function burden | ANOVA across subtypes | P = 0.30 |
| ALS gene panel carrier rate (34 genes) | chi-square | P = 0.95 (242 of 360 carriers, 67.2%) |
| ALS gene panel burden | Kruskal-Wallis | P = 0.98 |
| CCT2 rare damaging variants | carrier count | 0 of 360 patients |
| CCT2 cis-eQTL | linear model, 3,782 variants | minimum FDR q = 0.712 |
| CCT2 cis-eQTL in ALS GWAS | lookup, top 50 variants | 42 of 50 matched, minimum P = 0.116 |
| TRiC complex carrier rate | chi-square | P = 0.19 |
| TRiC complex burden | ANOVA | P = 0.35 |
| MYC target pathway burden | Kruskal-Wallis | P = 0.85 |
| MYC target carrier rate | chi-square | P = 0.73 |
| MYC target per-gene scan | Fisher, 128 genes | top hit XPOT, q = 0.362 |
| Genomic k-means vs transcriptomic subtype | adjusted Rand index | -0.005 |
| Blind genomic consensus clustering, k = 2 | adjusted Rand index | 0.003 (359/1 split) |

Rare variants are those with minor allele frequency below 0.01. Analyses restricted to the S-ALS-3 versus S-ALS-2 comparison were performed in 280 of these patients.

Genome-wide rare damaging variant burden did not differ across subtypes by ANOVA (P = 0.864; means 2,182, 2,220, and 2,207 variants for S-ALS-1, S-ALS-2, and S-ALS-3). The non-parametric equivalent sat at the conventional threshold (Kruskal-Wallis P = 0.05), which we report rather than selecting the more favorable test; the two disagree only marginally and the means are within 2% of each other. Rare loss-of-function burden did not differ either (ANOVA P = 0.30) (Fig. 7c). Carriage of rare damaging variants in a curated panel of 34 established ALS genes was common in this cohort (242 of 360 patients, 67.2%) but unrelated to subtype: carrier rates were 68.8%, 66.7% and 67.0% in S-ALS-1, S-ALS-2 and S-ALS-3 (chi-square P = 0.95), and the per-patient count of such variants did not differ either (Kruskal-Wallis P = 0.98). Variants were found in 33 of the 34 genes, with NEFH the most frequently affected (39.4% of patients).

Unsupervised clustering on genomic principal components did not recover the transcriptomic labels (adjusted Rand index = −0.005). Blind consensus clustering of the same genomic features produced only degenerate solutions: the k = 2 split isolated a single high-burden outlier from the remaining 359 patients, and its agreement with the transcriptomic partition was correspondingly negligible (adjusted Rand index = 0.003) (Fig. 7d).

At the CCT2 locus specifically, no rare damaging coding variant was observed in any of the 360 patients, including the 280 patients in the pre-specified S-ALS-3 versus S-ALS-2 comparison. This is therefore an absence of variants rather than a null association, since no burden test is estimable (Fig. 7a). There was no significant local cis-eQTL within 1 megabase of CCT2 (3,782 variants tested in 360 patients, minimum FDR q = 0.712; strongest variant chr12:70125111, P = 0.002) (Fig. 7b). Extending this, 42 of the 50 most strongly associated cis variants were present in the van Rheenen et al. (51) ALS genome-wide association study, and none showed even nominal association with ALS risk (minimum P = 0.116), arguing against a common regulatory variant linking CCT2 to disease susceptibility. Because no cis-eQTL survived correction in the first place, this is a lookup on nominal signals and does not exclude a regulatory variant undetectable at this sample size.

Burden across the eight-subunit TRiC complex did not differ by subtype (carrier chi-square P = 0.19; burden ANOVA P = 0.35), nor did burden across a 200-gene MYC target network (Kruskal-Wallis P = 0.85; carrier chi-square P = 0.73). A per-gene scan across the 128 MYC target genes with at least one carrier found a nominal enrichment of rare variants in XPOT, a tRNA export factor, in S-ALS-3 (5 of 88 patients versus 0 of 192 in S-ALS-2, Fisher P = 0.003), but this did not survive multiple testing correction (FDR q = 0.362) and we treat it as hypothesis generating only.

S-ALS-3 carried the lowest rare variant rate of the three subtypes in several overlapping tests (TRiC carriers 5.7% versus 12.5% in S-ALS-2; MYC target loss-of-function carriers 2.3% versus 7.3%), and every corresponding logistic coefficient pointed in the same direction at P values between 0.10 and 0.23. None is significant, but the direction is consistent. Rare variant burden across MYC target genes showed a nominal positive association with CCT2 expression (beta = 0.016, P = 0.006) that did not survive adjustment for genome-wide rare damaging burden (beta = 0.011, P = 0.066), with an effect roughly an order of magnitude smaller than the subtype difference. We report it as a disclosed non-finding.

The S-ALS-3 and CCT2 state therefore looks post-genomic: a regulatory state, not something written into the germline.

### The S-ALS-3/CCT2 state associates with markers of axonal injury and functional decline

We tested the clinical relevance of the framework against plasma neurofilament light chain (NfL, a marker of axonal damage), longitudinal ALSFRS-R decline slope, and disease duration.

For log NfL (n = 297), CCT2 expression (per standard deviation) was associated with higher NfL after adjustment for age, sex, and surrogate variables (beta = +0.039, P = 0.045), having been non-significant unadjusted (beta = +0.025, P = 0.177). Adding subtype membership to the model collapsed the CCT2 coefficient almost to zero (beta = +0.008, P = 0.752). Subtype membership was itself associated with NfL (omnibus F test P = 0.015), but neither term remained significant when both were in the model (subtype P = 0.11; CCT2 P = 0.75). Because CCT2 largely defines the S-ALS-3 state, the two terms are strongly collinear, and the association with NfL cannot be assigned to one or the other. We did not perform a formal mediation analysis.

ALSFRS-R decline followed a different pattern (n = 244). CCT2 expression (per standard deviation; slope in ALSFRS-R points per month) predicted faster decline both unadjusted (beta = −0.135, P = 0.002) and adjusted for age, sex, and surrogate variables (beta = −0.137, P = 0.005), and remained a similarly sized, still-significant predictor after adding subtype membership (beta = −0.122, P = 0.039), while subtype itself was not an independent predictor once CCT2 was in the model. Because differentiation quality was higher in S-ALS-3 than in S-ALS-2 (Table 2), we repeated this model with the proliferation, maturity and differentiation quality composites as additional covariates. The CCT2 coefficient fell by about a third and was no longer significant (beta = −0.084, P = 0.21), and adding the astrocyte, microglia and oligodendrocyte scores gave a similar result (beta = −0.076, P = 0.27). We therefore cannot exclude that part of the ALSFRS-R association reflects culture composition. In the models without composition covariates, CCT2 as a continuous graded predictor was the best-supported encoding of this endpoint by Bayesian information criterion (delta BIC 6.0 against the axis score and 9.0 against subtype). For NfL, by contrast, the CCT2-only and subtype-only models were statistically indistinguishable (delta BIC 1.2), so neither model comparison nor the nested models separate the gene from the state for this endpoint.

Disease duration could be computed for the 182 patients with both symptom onset and death recorded. It did not differ significantly across subtypes (ANOVA P = 0.20; Kruskal-Wallis P = 0.08; means 3.5, 4.2, and 3.5 years for S-ALS-1, S-ALS-2, and S-ALS-3), and CCT2 was null at every specification (beta = −0.469, P = 0.632 unadjusted). Missingness of the death date was not subtype-dependent (chi-square P = 0.57). Two limitations apply: duration is computed only among deceased patients, so it conditions on death and is not a survival analysis, and the parametric and non-parametric tests give different P values (0.20 and 0.08), although neither reaches 0.05. Age at death did not differ across subtypes after adjustment for onset age and sex (both P > 0.37). As external context, plasma NfL was a strong independent predictor of disease duration in this cohort (beta = −4.61 years per log10 NfL unit, P = 1.2 × 10^−9^, n = 142), supporting NfL as a valid severity marker generally, even though the subtype to duration link could not be tested directly.

The two endpoints run on separate tracks. For NfL, the CCT2 association cannot be separated from subtype membership. CCT2 itself tracks how fast patients lose function independently of the label, although this association weakens once culture composition is accounted for (Fig. 8c).

**Figure 8.**
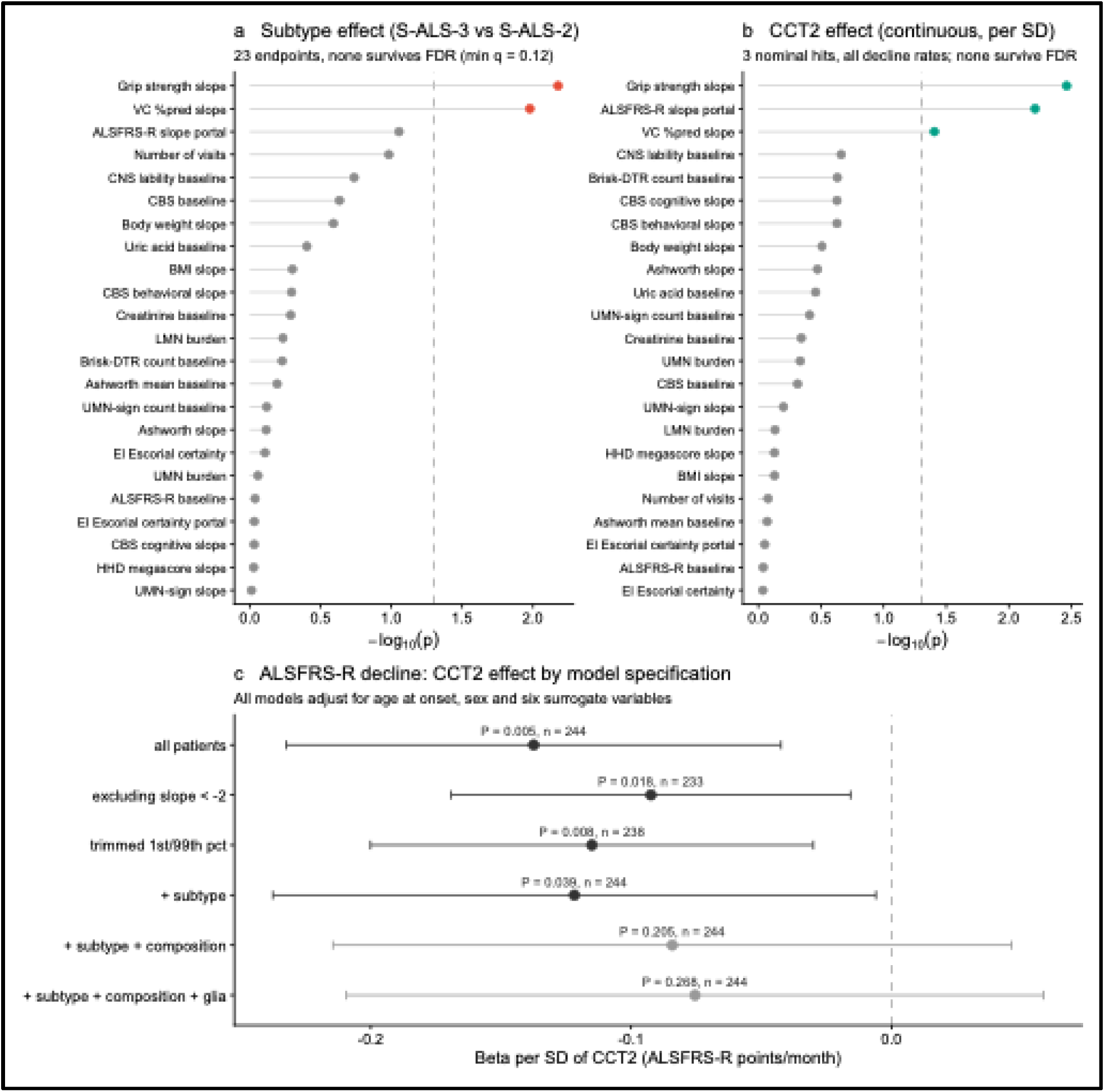
Clinical endpoint screen. (a) Subtype effect across the 23 modeled clinical endpoints, ordered by P value. Twenty-five endpoints were screened; two with fewer than 20 evaluable patients were not modeled. Benjamini-Hochberg correction was applied across the 23 modeled endpoints and no endpoint survives it (minimum q = 0.12). (b) CCT2 effect across the same endpoints, tested per standard deviation of expression. The three smallest P values are all rate-of-decline measures: grip strength slope, ALSFRS-R slope, and vital capacity slope. None survives correction across the 23 modeled endpoints. (c) The CCT2 association with ALSFRS-R decline (per standard deviation of CCT2) under six model specifications, all adjusted for age at onset, sex and six surrogate variables, showing the coefficient with 95% confidence interval. The effect holds when fast progressors are excluded, the distribution is trimmed, or subtype is added, and weakens when culture composition covariates are added. VC %pred, vital capacity percent predicted; CBS, ALS Cognitive Behavioral Screen; HHD, hand-held dynamometry; UMN and LMN, upper and lower motor neuron; DTR, deep tendon reflexes; CNS lability, Center for Neurologic Study lability scale; portal, AnswerALS portal-curated value.

### The progression signal is not explained by axonal injury alone

Because NfL is itself a strong predictor of ALSFRS-R decline in this cohort, we asked whether the transcriptomic predictors carry progression information beyond it. Adding NfL to the ALSFRS-R model raised adjusted R squared from 0.058 to 0.262 (F = 49.3, P = 4.8 × 10^−11^, n = 188), confirming NfL as the dominant single predictor of decline rate.

Against that background, the metabolic axis and the discrete subtype label did not survive. The metabolic axis was marginal alone (beta = −0.085, P = 0.092) and attenuated with NfL in the model (beta = −0.060, P = 0.183); the S-ALS-3 label was non-significant alone (beta = −0.202, P = 0.110) and collapsed with NfL added (beta = −0.037, P = 0.750, n = 148). CCT2 expression (per standard deviation) behaved differently: it predicted faster decline alone (beta = −0.142, P = 0.010) and remained significant with NfL in the model (beta = −0.099, P = 0.047), retaining roughly 70% of its coefficient. The effect is modest and the attenuation is real, but CCT2 carries progression-rate information that axonal injury does not fully explain. The discrete subtype label does not. No individual Hallmark pathway was independently associated with any clinical endpoint across 30 pathway-by-endpoint combinations (smallest FDR = 0.213; Supplementary Note 3).

### An extended clinical endpoint screen points to a progression-specific pattern

We screened 25 candidate clinical endpoints, of which 23 had at least 20 evaluable patients and were modeled for association with subtype membership and with CCT2 expression (Fig. 8a-b).

No endpoint separated the subtypes after correction for multiple testing. The two strongest were grip strength decline (S-ALS-3 beta = −15.8 lb/yr, P = 0.007, FDR = 0.121, n = 66) and vital capacity percent-predicted decline (beta = −9.5 %pred/yr, P = 0.011, FDR = 0.121, n = 163); all other endpoints had larger P values still.

CCT2 expression (per standard deviation, as in Fig. 8b) showed a more focused pattern. The three lowest CCT2 P values across the 23 modeled endpoints were all rate-of-decline measures: grip strength slope (beta = −9.9 lb/yr, P = 0.003, FDR = 0.071), portal-curated ALSFRS-R slope (beta = −0.153 /mo, P = 0.006, FDR = 0.071), and vital capacity slope (beta = −4.0 %pred/yr, P = 0.039, FDR = 0.301). None survives correction across the 23 modeled endpoints, and we do not claim any single one as significant. The reportable observation is the pattern: the discrete subtype label separates nothing after correction, while CCT2 concentrates its nominal associations specifically in measures of functional decline rate.

All remaining endpoints were null for both predictors (Supplementary Table S2). Cohort retention, measured as number of study visits, did not differ by subtype (P = 0.104), so there is no sign of differential dropout between subtypes.

CCT2 picks out decline rate and nothing else in this screen. It does not track baseline severity, motor neuron burden, spasticity, diagnostic certainty, or cognition, all of which are conventional ways of stratifying ALS patients. Whatever it is measuring is close to orthogonal to them. One caveat applies across every longitudinal slope endpoint in this screen: patients who decline fastest are also the least likely to remain in the cohort long enough to accumulate the visits required for a slope estimate, so all slope-based results here carry some risk of survivor bias in the same direction.

### Cross-omic and cross-tissue comparisons support the direction of the familial-sporadic geometry

We compared C9orf72 and sporadic ALS in an independent laser-capture microdissected postmortem motor neuron dataset (GSE56500; 3 C9orf72, 3 sporadic, and 6 control subjects) (52). Gene-level fold changes correlated at Spearman rho = 0.59, and Hallmark enrichment scores at rho = 0.96 across all 50 pathways, with 23 of the pathways significant in either contrast significant in both (Jaccard 0.85, Fisher P = 5.2 × 10^−10^). The direction agrees with our iPSC-MN finding, but the magnitudes should not be compared directly: with three subjects per disease group, both contrasts are dominated by a shared end-stage degenerative signal, and cross-genotype correlations in postmortem tissue are systematically inflated relative to purified motor neurons, as the Target ALS bulk comparison above demonstrates (rho = 0.75). The microdissected estimate (0.59) falls between our purified iPSC-MN estimate (0.33) and the bulk postmortem estimate (0.75), consistent with cell type purity rather than disease stage driving the difference. We read this as directional corroboration, not replication.

We also compared our subtypes against Tam et al. (42), who described three postmortem transcriptomic ALS subtypes, ALS-Glia, ALS-Ox, and ALS-TE, in bulk cortical and spinal cord tissue. Correspondence was partial: S-ALS-1, our inflammatory, pan-activated state, shares thematic overlap with ALS-Glia, and S-ALS-3, our metabolically dysregulated state, shares thematic overlap with ALS-Ox. Neither S-ALS-2 nor ALS-TE has a clear analog in the other dataset, which is expected given differences in tissue (purified motor neurons versus bulk postmortem tissue dominated by glia) and disease stage (early to mid-course living patients versus end-stage autopsy tissue). This is context for plausibility, not cross-cohort validation.

### Spatial transcriptomics supports the CCT2 to cell cycle relationship in an independent cortical cohort

We examined CCT2 and the three axes in postmortem cortex from the Target ALS Postmortem Tissue Core, using 10x Visium spatial transcriptomics. Samples were pseudobulked from spots enriched for layer 5b projection neuron markers (Methods). This yielded 28 ALS and 11 control frontal cortex profiles (Fig. 9).

**Figure 9.**
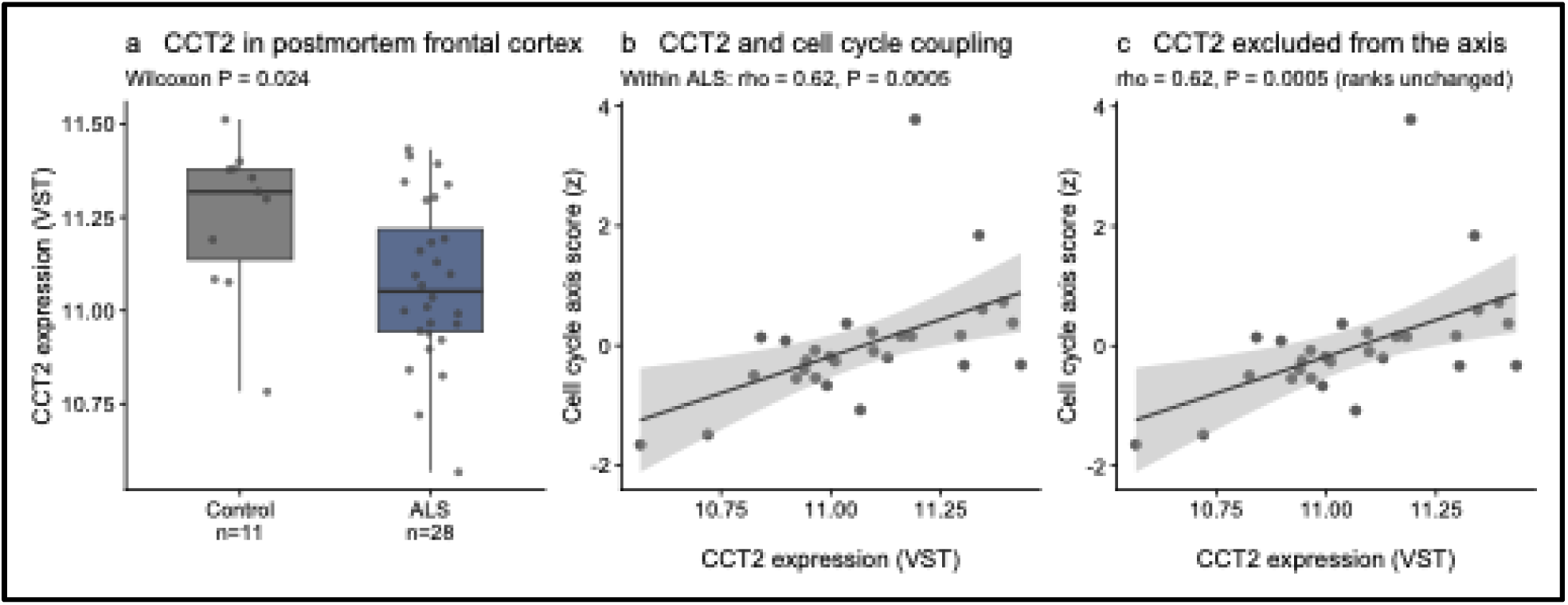
Spatial transcriptomic validation in postmortem frontal cortex. (a) CCT2 expression in ALS and control frontal cortex from the Target ALS Postmortem Tissue Core. Each point is one subject, pseudobulked from the top quartile of Visium spots by a layer 5b projection neuron marker score. (b) CCT2 against the cell cycle axis score within the ALS group; the line is a linear fit with 95% confidence band. (c) The same relationship with CCT2 removed from the cell cycle gene set. Sample ranks are unchanged and the rank correlation is identical, so CCT2 contributes negligibly to the composite score.

Between groups, CCT2 was lower in ALS than in control (Wilcoxon P = 0.024). Two factors likely contribute. In the iPSC-MN cohort, CCT2 is below control in S-ALS-1 and S-ALS-2, which together make up about three quarters of sporadic patients, so a lower ALS group mean is expected even without cell loss. End-stage cortex has also lost projection neurons, in which CCT2 is highly expressed. A group comparison in postmortem tissue cannot separate these two, and it cannot show the S-ALS-3 minority moving in the opposite direction (Fig. 9a).

The within-ALS relationship is the more informative test. Among the 28 ALS samples, CCT2 expression correlated specifically with the cell cycle axis (Spearman rho = 0.62, P = 5.2 × 10^−4^; Pearson r = 0.53, P = 0.004), mirroring the axis-specific relationship seen in the primary iPSC-MN cohort, and not with the metabolic axis (rho = 0.15, P = 0.43) (Fig. 9b). The correlation was rho = 0.70 (P < 0.001) after removing the patient with the highest CCT2 and rho = 0.64 (P < 0.001) after removing the three highest. A nominal negative correlation with the inflammation axis (rho = −0.38, P = 0.045) did not survive correction across the three axes tested, and the inflammation axis itself differed between ALS and control samples (P = 0.040).

Because CCT2 is itself a member of the cell cycle gene set, we recomputed the axis score with CCT2 excluded. Sample ranks were unchanged and the rank correlation was identical (rho = 0.62), so CCT2 contributes negligibly to the composite score and the correlation is not circular (Fig. 9c).

The CCT2 to cell cycle relationship therefore holds in a different tissue, a different platform, and a different cohort. The discrete three-subtype partition does not travel with it; testing that would need subtype-annotated postmortem data, which does not currently exist.

## Materials and Methods

### Data access and cohort

We requested data from the AnswerALS (49) consortium and the request was granted on 18 April 2025. We downloaded the modalities used here: whole genome sequencing level four data (joint per-chromosome patient VCFs), proteomics level four data, transcriptomics level four data, and clinical data from the Neuromine portal. For the familial versus sporadic comparison, we used the genetic mutation table in the clinical data to identify patients with familial mutations; patients without any of the tested mutations were classified as sporadic.

The dataset comprises 644 RNA-seq samples from 619 participants: 400 samples from 391 sporadic patients, 46 from 43 C9orf72 carriers, 13 from 12 SOD1 carriers, and 185 from 173 controls. An earlier assembly of this cohort treated repeated sequencing of a single reference differentiation control line as 98 independent control participants. This was found during quality control and every analysis reported here was recomputed on the corrected cohort.

Two analytic levels are used throughout and are distinguished explicitly. Differential expression, surrogate variable estimation, ssGSEA scoring and consensus clustering were performed at the sample level. Each patient’s subtype label was taken from the earliest available sample, and classifier training and all clinical and genomic association analyses were performed at the patient level. Where sample-level and patient-level counts differ, both are given. The cohort was roughly balanced by sex overall, though the sporadic group skewed male and the control group skewed female, and age at symptom onset peaked between 50 and 60 years (Fig. 1).

### Transcriptomic processing and differential expression

RNA sequencing was processed through a standard filtering and normalization pipeline, retaining 24,253 genes after requiring at least 10 counts in at least 10% of samples, of which 18,606 mapped to unique gene symbols and formed the matrix used for all gene set scoring. Unmodeled technical structure was captured with surrogate variable analysis (SVA), using two separate fits described in the next paragraph (53). Differential expression was then fitted as ∼ sex + the six genotype-independent SVs + mutation subtype, where mutation subtype is the grouping variable that defines each contrast rather than a covariate. A parallel proteomics dataset (436 participants, 3,256 quantified proteins) was used to test whether transcriptomic findings replicated at the protein level.

Two SVA fits were used, and they are not interchangeable. The backbone fit, used for axis scoring, clustering and all analyses within the sporadic cohort, was run on the full 742-sample set including the reference differentiation line, which contributes a strong and well-defined technical signal. The resulting variables were applied to the corrected cohort, and the reference line was excluded from all cohort counts and contrasts. Axis construction and clustering used sex together with the first six backbone SVs (SV1 to SV6). For this purpose, refitting SVA on the corrected cohort alone was tested and rejected, because the resulting surrogate variables correlated 0.50 to 0.65 with the biological axes and removed the subtype structure along with the technical structure. The genotype fit, used only for the familial versus sporadic contrasts, was run on the 644-sample corrected cohort. Here the concern was different: surrogate variables that track genotype would absorb the contrast being tested. Each of the ten variables recovered was tested for association with mutation subtype by one-way ANOVA, and the three with P < 0.05 (SV3, SV4 and SV5; P = 0.004, 0.022 and 0.023) were excluded. Of the seven that remained, the first six in index order (SV1, SV2, SV6, SV7, SV8 and SV9) were retained, so that the genotype fit uses the same number of surrogate variables as the backbone fit. SV10 was not associated with genotype (P = 0.25).

Genotype proximity was assessed at gene and pathway level. At the gene level we computed the Spearman correlation of log2 fold changes, and directional agreement on the top 500 genes, defined both as the top 500 in each pairwise comparison and, separately, as a single fixed reference set of the 500 genes most significant in C9orf72 evaluated in the other groups. At the pathway level we used the Spearman correlation of normalized enrichment scores across all 50 Hallmark gene sets. The same comparisons were run on the proteomics data.

### Proteomics

Proteomic data were generated by the AnswerALS consortium and processing is described in full in the consortium’s methods (49). In brief, peptides were acquired on a 6600 Triple TOF (Sciex) in both data-dependent mode, to build a sample-specific ion library, and data-independent mode over 100 variable windows; DIA files were quantified against that library with OpenSWATH and protein-level differential abundance computed with mapDIA. After mapping to gene symbols, 3,256 proteins across 436 participants were available here. CCT2 is absent from the symbol-collapsed matrix and was recovered from the pre-mapping protein matrix under its UniProt identifier (P78371, TCPB_HUMAN); this step is required to reproduce the CCT2 protein results.

### Postmortem comparison datasets

Bulk RNA-seq from postmortem motor cortex was obtained from the Target ALS Postmortem Tissue Core, and laser-microdissected motor neuron data from a GEO study (52), GSE56500, comprising 3 C9orf72, 3 sporadic and 6 control subjects. For the Target ALS bulk data, we retained motor cortex lateral and medial samples, assigned genotype from the C9orf72 repeat expansion field together with string matching on reported genomic mutations, and fitted DESeq2 with sex and group, giving 43 control, 53 C9orf72, 18 SOD1 and 320 sporadic samples (54). For GSE56500 we fitted limma on the processed series matrix (55). Given the sample size, the GSE56500 comparison is reported as directional only.

### Axis definition

From the Hallmark pathway analysis, we derived the recurring pathways and pre-specified three axes before any clustering: AXIS_METABOLIC (oxidative phosphorylation, fatty acid metabolism, glycolysis; 512 genes), AXIS_CELL_CYCLE (E2F targets, G2M checkpoint, MYC targets V1, DNA repair; 610 genes) and AXIS_INFLAMMATION (TNF alpha signaling via NF-kB, interferon gamma response, complement; 538 genes) (Supplementary Table S1). Each patient received one ssGSEA (56) score per axis, computed on the sex- and surrogate-variable-residualized expression matrix and z-scored across patients.

To test whether the three-axis structure depended on the Hallmark database, we rebuilt all three axes from GO Biological Process (MSigDB C5) and Reactome (C2:CP:REACTOME) gene sets via msigdbr, rescored all patients on the same residualized matrix, and compared both the axis scores and the downstream cluster assignments against the Hallmark framework by adjusted Rand index, inspecting cluster sizes in every case to exclude degenerate partitions(50). As a further check we tested a fully data-driven alternative, submitting genes with adjusted P below 0.05 and absolute log2 fold change above 0.3 in each of the C9orf72, SOD1 and sporadic versus control contrasts to GO Biological Process over-representation analysis (clusterProfiler, Benjamini-Hochberg correction, all tested genes as background)(57).

### Consensus clustering

We ran ConsensusClusterPlus (58) on the three axis scores for the 400 sporadic samples, across k = 2 to 5 (1,000 resamples, 80% item sampling, hierarchical clustering, Euclidean distance, seed 42), choosing k by minimizing the proportion of ambiguous clustering. Resulting clusters were compared against the axes to identify which cluster aligned with each. A second round of clustering on expression residualized against measured differentiation marker percentages was run and rejected: it produced a partition with a single-patient cluster, which cannot be meaningfully compared with any other. Only the first round is reported.

### Cell composition and differentiation quality

Composite scores were the mean of row-z-scored expression across each marker panel. Differentiation quality was the maturity composite (14 markers including MAP2, SYN1, NEFL, NEFM, SNAP25, RBFOX3, STMN2, CHAT, ISL1, MNX1) minus the progenitor composite (9 markers including PAX6, SOX2, NES, MKI67, TOP2A, PCNA). Panel completeness in this dataset was: maturity 14 of 14, progenitor 9 of 9, proliferation 7 of 7, astrocyte 7 of 8 (GFAP absent), oligodendrocyte 5 of 6 (MOG absent) and microglia 2 of 7 (AIF1, CX3CR1, P2RY12, CD68, TYROBP absent). The sparse microglial panel is expected in an iPSC-derived motor neuron culture, which contains no microglia, but the microglial adjustment therefore rests on two genes and is weak.

Three sensitivity analyses of increasing stringency were performed. First, we recomputed axis scores after regressing out S100B and re-clustered on the residuals. Second, we tested the differentiation quality composite across subtypes by one-way ANOVA with eta squared and Tukey post-hoc comparisons. Third, we regressed each axis score on the available astrocyte (7 of 8), microglia (2 of 7) and oligodendrocyte (5 of 6) markers together with the differentiation quality composite and re-clustered on the residuals. Sensitivity re-clustering used k-means consensus clustering (maxK 3, 1,000 resamples, 80% item sampling, Euclidean distance, seed 42) whereas the primary partition used hierarchical clustering, so the adjusted Rand indices reflect a change of algorithm as well as of input and are conservative in that respect.

### Classifier training and CCT2 characterization

We trained one-vs-rest binary LASSO classifiers on the 1,470 Hallmark axis genes present in the expression matrix for the 391 sporadic patients, with nested 10-by-10 cross-validation and regularization at lambda.1se optimized on AUC. Feature stability was assessed by bootstrap resampling. Classifier performance is reported at the patient level; the ROC curve for CCT2 alone was computed from ranks with a stratified bootstrap confidence interval.

CCT2 expression across clusters was compared by mean VST, one-way ANOVA with effect size, and Tukey post-hoc correction. To confirm that elevated CCT2 was a property of the S-ALS-3 state rather than an artifact of distributional overlap, multimodality or a few high-expressing samples, we applied three checks to the VST values: the number of S-ALS-3 patients exceeding the maximum S-ALS-2 value; Hartigan’s dip test for unimodality within S-ALS-3 (diptest); and repetition of the S-ALS-3 versus S-ALS-2 comparison after excluding the 15 S-ALS-3 patients with the highest CCT2.

### Upstream regulator and pathway analysis

Differential expression between S-ALS-3 and S-ALS-2 was performed with DESeq2 (54) on 311 samples (107 S-ALS-3, 204 S-ALS-2), using the design ∼ sex + backbone SV1 to SV6 + subtype, with S-ALS-2 as the reference level. The full result table, 24,021 genes after excluding sex chromosome genes, was uploaded to Ingenuity Pathway Analysis (QIAGEN, version 01-23-01). Genes with a false discovery rate below 0.001 and an absolute log2 fold change of at least 0.3 were selected for analysis (2,865 genes; 771 higher and 2,094 lower in S-ALS-3) and analyzed by Core Analysis against the Ingenuity Knowledge Base (Genes only), using direct and indirect relationships and experimentally observed confidence level. The analysis returned 3,882 upstream regulators and 1,187 canonical pathways, reported with their activation z-score and overlap P value. As an independent check, GO Biological Process over-representation analysis was run with clusterProfiler (version 4.18.4) separately on the genes higher and lower in S-ALS-3 from the same list, against all tested genes as background, with Benjamini-Hochberg correction, and redundant terms were collapsed with simplify()(57). To locate the source of the cell cycle axis difference, the four Hallmark sets making up that axis were scored separately by ssGSEA (GSVA version 2.4.9) on the unresidualized VST matrix and compared between S-ALS-3 and S-ALS-2 patients by Wilcoxon rank-sum test(59).

### Germline genomic analyses

We ran a series of tests at the WGS level to ask whether the transcriptomic subtypes are visible in inherited DNA, using whole genome sequencing matched to RNA-seq, available for 360 sporadic patients. All genomic analyses were performed in this set. Analyses restricted to the pre-specified S-ALS-3 versus S-ALS-2 comparison used the 280 patients in those two subtypes. Rare variants were those with minor allele frequency below 0.01, limited to predicted-damaging coding variants, summarized per patient as counts of total damaging, loss-of-function and nonsynonymous variants from the AnswerALS ANNOVAR reports(60).

Two variant-parsing conventions are present in the pipeline and are noted for transparency: the genome-wide burden counts a carrier when the genotype is 0/1, 1/0 or 1/1, whereas the TRiC and MYC target burden counts any alternate allele, including multi-allelic genotypes such as 1/2. Results are null under both.

i. Rare-variant burden was tested for association with subtype membership genome-wide by one-way ANOVA and Kruskal-Wallis, and within a curated panel of 34 established ALS genes by chi-square test of carrier rates and Kruskal-Wallis test of per-patient variant counts (49).
ii. Unsupervised genomic clustering. We clustered patients on genomic principal components by k-means and, separately, by blind consensus clustering, comparing each against the transcriptomic labels by adjusted Rand index and inspecting cluster sizes.
iii. CCT2 local regulation. We ran a cis-eQTL analysis for all common variants (MAF at least 0.01; 3,782 variants) within a 1 Mb window centered on CCT2 (chr12:69.2 to 70.3 Mb, GRCh38), with Benjamini-Hochberg correction across tested variants. Variants were extracted and additively dosage-coded with PLINK (--recode A) and tested by per-variant linear regression adjusting for sex and the six genotype-independent SVs, with one observation per participant(61). Top cis variants were then looked up in the van Rheenen et al. (51) ALS GWAS after aligning both datasets to GRCh38 using the GWAS Catalog harmonized release of that study (accession GCST90027164); harmonized effect alleles and effect sizes were used throughout and allele orientation was checked per variant before any direction-of-effect comparison.
iv. CCT2-network rare-variant burden was tested at three nested scopes: CCT2 alone, the eight-subunit TRiC/CCT chaperonin complex (pooled chi-square, plus negative binomial and logistic regression adjusting for age and sex), and a 200-gene MYC target network, each tested against subtype membership. Because pathway-level burden counts partly index each patient’s global rare-variant load, the MYC target association with CCT2 expression was additionally adjusted for genome-wide rare damaging burden.

Pathway-specific polygenic scores were built with PRSet (PRSice v2.3.5) from the van Rheenen et al. (51) summary statistics and are not reported. The summary statistics used were the GRCh37 release of that study while the AnswerALS genotypes are on GRCh38, so fewer than 1% of variants matched by position and the resulting scores could not be interpreted. Repeating this analysis on build-matched summary statistics is straightforward and is left to future work.

### Clinical associations

Clinical associations were tested for plasma neurofilament light chain (NfL), ALSFRS-R progression slope and disease duration. Plasma NfL was log10 transformed. For each endpoint, associations with subtype membership were assessed by linear regression adjusting for age at symptom onset, sex and the six backbone SVs, evaluated for the pre-specified S-ALS-3 versus S-ALS-2 contrast. Continuous transcriptomic predictors (the metabolic axis score and CCT2 expression) were tested against each endpoint with the same covariate-adjusted models. Because differentiation quality differed between subtypes, the CCT2 models for ALSFRS-R and NfL were repeated with the proliferation, maturity and differentiation quality composites as additional covariates, and then with the astrocyte, microglia and oligodendrocyte scores added. Robustness of the ALSFRS-R association was also checked by excluding patients with slopes below −2 points per month and by trimming slopes at the 1st and 99th percentiles.

To distinguish whether CCT2’s relationship to each endpoint operated through subtype membership or independently of it, we compared nested models per endpoint: endpoint ∼ CCT2 + covariates, and endpoint ∼ CCT2 + subtype + covariates. Attenuation of the CCT2 coefficient on adding the subtype term was read as indicating that the association could not be separated from the categorical state; stability of the coefficient as a subtype-independent graded association. Model comparison used the Bayesian information criterion. Because NfL is itself a strong predictor of ALSFRS-R decline, each transcriptomic predictor was also tested with NfL added to the model.

Disease duration was computed as age at death minus age at symptom onset and is available only for deceased patients. It therefore conditions on death and is not a survival analysis; missingness was tested for dependence on subtype.

To test whether any single pathway component could be identified as a driver of outcome, we scored each sporadic patient on the 10 individual Hallmark gene sets underlying the three axes by ssGSEA, z-scored across patients, and fit endpoint ∼ pathway + age at onset + sex + the six backbone SVs for every pathway by endpoint combination, using log NfL (n = 297), ALSFRS-R slope (n = 244) and cognitive progression slope (n = 135). This gave 30 marginal models corrected jointly by Benjamini-Hochberg, with joint models including all 10 pathways fit per endpoint to account for collinearity.

To determine whether the subtype and CCT2 associations extended to other domains of ALS assessment, we screened every quantifiable measure available in the AnswerALS clinical forms and data portal against subtype (S-ALS-2 as comparator, S-ALS-3 as the group of interest) and CCT2 expression, giving 25 endpoints of which 23 had at least 20 evaluable patients and were modeled. Linear endpoints used endpoint ∼ subtype + age at onset + sex + the six backbone SVs, with CCT2 tested univariately and then alongside subtype. Benjamini-Hochberg correction was applied across the screen. Longitudinal slope endpoints required at least three visits spanning at least 90 days; patients with a single visit were dropped rather than zero-imputed.

### Spatial transcriptomics

To test whether the CCT2 to cell cycle relationship generalizes beyond the iPSC-MN system, we analyzed an independent spatial transcriptomic dataset identified through the Target ALS Data Engine: Visium spatial RNA-seq generated by Columbia University on postmortem cortex, spanning 43 subjects (30 ALS, 13 non-neurological control) across frontal cortex (39 subjects) and motor cortex (4 subjects, split medial and lateral). The Target ALS Postmortem Tissue Core banks frozen and FFPE tissue from 300 ALS and 50 control cases across five US sites and distributes histopathology, whole genome sequencing, bulk RNA-seq and spatial transcriptomic data to approved investigators; access was granted following a short research proposal. Not every data type exists for every case.

Given the small motor cortex n, we treated the frontal cortex arm as the primary test (28 ALS, 11 control) and the motor cortex arm as a descriptive case series only. Raw and processed sequencing outputs and histopathology images were retrieved through the Data Engine file manifests, filtered to the minimal file set needed (filtered feature-barcode matrix, spot coordinates, scale factors and the high-resolution tissue image) rather than the full Space Ranger bundle.

For each capture area, spots enriched for layer 5b cortical projection neurons were identified with a published marker panel (FEZF2 and CRYM for general layer 5b identity; BCL11B, POU3F1 and NEFH for corticospinal and Betz cell enrichment (62)), scored per spot with Seurat (63)module scoring, and the top quartile of spots by score was pseudobulked per subject and tissue. Each sample is therefore enriched for layer 5b projection neurons rather than being a whole-section average. Because the primary arm is frontal cortex, which lacks Betz cells, the corticospinal markers index layer 5b projection neurons generally. These are cortical neurons, not the spinal motor neurons that iPSC-MN differentiation models, so this analysis tests whether the CCT2 to cell cycle relationship generalizes to another neuronal population rather than replicating the same cells. Pseudobulk profiles were normalized with DESeq2 VST (54), and the identical AXIS_METABOLIC, AXIS_CELL_CYCLE and AXIS_INFLAMMATION definitions and ssGSEA procedure used in the primary analysis were applied unchanged.

Group differences in CCT2 between ALS and control were tested by two-sided Wilcoxon rank-sum test. Within the ALS group, the relationship between CCT2 and each axis score was tested by Pearson and Spearman correlation, and sensitivity to high-leverage points assessed by sequentially removing the one and three patients with the highest CCT2 and repeating the correlation. Because CCT2 is a member of the cell cycle gene set, the axis was also recomputed with CCT2 removed and the correlation repeated.

## Discussion

Sporadic ALS accounts for roughly 90% of cases, yet unlike familial disease it offers no shared genetic anchor, and its biological heterogeneity is a recognized reason that trials enrolling patients under a single diagnosis have struggled to show treatment effects (29,33–35). Hence for sporadic ALS, stratification has to come from the biology itself. Here we used iPSC-derived motor neurons from 391 living sporadic patients in the AnswerALS cohort to ask whether that biology is structured. We found that the transcriptional programs separating patients organize along three axes, metabolic, cell cycle, and inflammatory, and that patients fall into three states along them. One of these, S-ALS-3, can be identified by a single gene, CCT2, whose expression is associated with ALSFRS-R decline, although this association weakens after adjustment for culture composition. Previous efforts to subtype ALS have relied mainly on postmortem tissue (39–42). That tissue captures end-stage disease and is dominated by glia and infiltrating immune cells, which, as our Target ALS and GSE56500 comparisons show, compresses the differences between genotypes. Patient-derived motor neurons trade the tissue environment for cell-type purity and an earlier point in the disease course, which is the setting in which stratification would actually be applied.

Prior subtyping studies conducted on postmortem tissue have described three main subtypes, ALS-Ox, ALS-Glia and ALS-TE (40–42). Comparison with our subtypes shows a partial overlap. S-ALS-1, which has high inflammation and is activated on all three axes, is closest to ALS-Glia, and S-ALS-3, which has high metabolic dysregulation, is closest to ALS-Ox. Neither ALS-TE nor S-ALS-2 has a counterpart in the other dataset. This pattern of partial overlap is expected. Postmortem tissue is end-stage and is mostly made up of glia, while iPSC motor neurons are a single cell type taken from living patients at an earlier point in the disease, and the two are not going to show the same subtypes. ALS-TE not having a counterpart in our data could be because retrotransposon activation has been linked to TDP-43 pathology, which iPSC motor neurons mostly do not develop in culture, and because our gene-level counts do not include transposable elements. S-ALS-2 not having a counterpart in postmortem data could be because it is the quiet subtype, low on all three axes, and there may be no place for such a subtype to show up in end-stage tissue where every sample has glial activation and neuronal loss. We also acknowledge a limitation of the AnswerALS cohort itself. A recent study pointed out that the transcriptomic, epigenetic and proteomic data from AnswerALS have not shown a clear separation between sporadic patients and controls, and that its differentiation protocol gives more variable cultures than an optimized protocol with over 92% motor neuron purity (47). Our study asks a different question, whether there is structure within sporadic ALS rather than whether sporadic ALS separates from controls, but the variable culture composition is still a concern for us, particularly for S-ALS-1, and we address this in the limitations below.

The main practical finding of this study is that S-ALS-3 can be identified with a single gene. CCT2 came out of a LASSO screen of over a thousand candidate genes and it was selected for S-ALS-3 only, not for the other two subtypes. The full LASSO panel does slightly better than CCT2 alone, so the case for a single gene is not accuracy. It is that one transcript or one protein is something that can be measured in a trial or a clinic, while a panel of twelve to sixteen genes is harder to carry across cohorts and platforms. A natural concern with a single-gene marker is that a few patients with very high expression are driving the separation. This is not the case here. The CCT2 distribution in S-ALS-3 is unimodal, and the separation from S-ALS-2 holds after the highest patients are removed. The signal is also present at the protein level in the subset of patients with proteomics, although the correlation between transcript and protein is modest and the proteomics depth in this cohort is limited. CCT2 is a subunit of the TRiC chaperonin, which folds actin, tubulin and a number of cell cycle proteins (64,65). It would be tempting to read this as TRiC dysfunction in S-ALS-3, but a composite score of all eight TRiC subunits does not separate the subtypes. What separates them is CCT2 specifically. We therefore treat CCT2 as a marker of a regulatory state in these motor neurons and not as evidence that the chaperonin itself is the driver. That leaves the question of what is pushing CCT2 up in this subset of patients. CCT2 has come up in ALS before, but not in this way. One study using this same cohort listed seven CCT subunits together as candidate drug targets (66). That list came from expanding a protein interaction network around genes picked by machine learning, so the whole complex came in as a block, and it was not a finding about patients. Our result points the other way. The TRiC complex as a whole does not separate our subtypes. CCT2 alone does, and that is what makes it useful as a marker.

CCT2 has also been studied in neurodegeneration from a different direction. It works as an aggrephagy receptor, and raising CCT2 expression clears aggregation-prone proteins including mutant SOD1 and FUS (67). Reduced CCT2 has been reported to track cellular injury in Alzheimer’s disease (68). Our data fit that picture for most patients. Compared with controls, CCT2 is lower in S-ALS-1 and S-ALS-2, which together are about three quarters of the sporadic cohort. S-ALS-3 is the exception and sits above control. So the disagreement is not between our study and the functional literature, it is between one subtype and the rest of the cohort. Because the two directions cancel, sporadic ALS as a whole is barely different from control on this gene, which is the same problem that makes unstratified comparisons in this cohort hard to interpret. Why S-ALS-3 goes the other way is not settled here. The elevation may be a response to proteostatic demand rather than a protective reserve, and in our data CCT2 sits inside a MYC and mTORC1 translation program rather than tracking the chaperonin complex, which does not separate the subtypes. We treat CCT2 as a marker of a regulatory state and not as a driver, so its direction does not have to match what overexpression does in a cell model. Whether CCT2 is doing chaperone work, aggrephagy work, or neither in S-ALS-3 will need experiments in the cells.

The transcriptional context of S-ALS-3 points to MYC and mTORC1. In the upstream regulator analysis, MYC and MYCN are predicted activated in this subtype, RPTOR is predicted activated, and the mTOR inhibitors torin1 and sirolimus are predicted inhibited, all of which describe the same thing from different sides, an active mTORC1. LARP1 is predicted inhibited as well. This is the expected pattern when mTORC1 is on, since mTORC1 phosphorylates LARP1 and that relieves its repression of the 5′TOP mRNAs (69,70). The LARP1 prediction needs one qualification. It is inferred from transcript abundance, whereas LARP1 acts mainly at the level of translation and also stabilizes TOP transcripts (71), so the two activities do not have to move transcript levels in the same direction. We treat it as consistent with 5′TOP de-repression rather than as demonstrating it. Polysome profiling in CCT2-high and CCT2-low lines would be the direct test. The relationship between CCT2 and this program may also run in both directions. CCT2 protein stabilizes c-Myc (72), and the CCT complex folds both mLST8 and Raptor, two components of the mTOR complexes (73). So elevated CCT2 in S-ALS-3 could be reinforcing the same growth signaling that raised it, rather than only reading out from it. We were careful about one more thing. CIP2A, which stabilizes MYC by inhibiting PP2A (74) and has been proposed as a link between MYC and mTORC1 in cancer (75), is not predicted activated in S-ALS-3. Whatever is holding MYC and mTORC1 active in these cells, it does not appear to be the CIP2A route. We also looked for a genetic explanation and did not find one, which we come back to below. What this program means for a post-mitotic motor neuron is not obvious. Cell cycle protein expression has been reported in ALS motor neurons at autopsy (76), and cell cycle re-entry has been linked to neuronal death across several neurodegenerative diseases (77). Our data argue against simple mitotic re-entry: the S-ALS-3 elevation on the cell cycle axis is carried by MYC targets, while G2M checkpoint genes are lower than in S-ALS-2. Whether any part of a re-entry program is active in S-ALS-3 cannot be settled from expression data and will need experiments in the cells themselves.

One of the results we consider most informative is a negative one. The subtypes are not explained by genetics. None of the sporadic patients with whole genome sequencing carry a damaging variant in CCT2, the CCT2 cis-eQTL scan is null, and the burden of rare damaging variants does not differ between subtypes, apart from one borderline non-parametric test that we report in full. Genomic clustering of the same patients bears no relationship to the transcriptional subtypes. If anything, S-ALS-3 has the lowest rare variant rate of the three, both in the TRiC genes and in MYC pathway genes. We think this is the right way to read the subtypes. They are regulatory states, set up downstream of whatever genetic and environmental exposures the patient carries, rather than genetic categories in disguise. This matters for how the subtypes might be used. A genetic subtype can be read off DNA once. A regulatory state has to be measured from the cells, and it may in principle be modifiable. It also fits with what we saw in the familial arm of the cohort, where C9orf72 and SOD1 carriers do not converge on a common transcriptional signature in this system. If two known mutations do not produce the same motor neuron state, it should not be surprising that sporadic patients without a shared mutation still organize into distinct states.

The clinical associations are modest and we present them as such. CCT2 expression is associated with faster decline on the ALSFRS-R, and this association holds after the subtype label is added to the model, so it is carried by the gene and not simply by membership in S-ALS-3. It does, however, lose about a third of its size and its significance when culture composition markers are added, so part of it may reflect how well the lines differentiated. CCT2 is also associated with plasma NfL, but here the picture is different. Once subtype is in the model the CCT2 term drops away, and model comparison cannot decide between the two, which means the NfL signal is shared between the gene and the subtype and we cannot say which is doing the work. Disease duration does not differ by subtype. In a screen of 23 modeled clinical endpoints, no subtype association survives correction for multiple testing, and the three endpoints where CCT2 is nominally associated, grip strength, ALSFRS-R and vital capacity, do not survive it either. We do not think this is a reason to dismiss the finding. The molecular data are from one sample per patient, the clinical data are available for different numbers of patients for different endpoints, and the cells were collected at a single visit. Eshima et al. found survival differences between their postmortem subtypes (40). The equivalent test in our cohort is underpowered, and we do not claim it. What we can say is that the one gene that defines S-ALS-3 is also the one that shows the most consistent, if weak, relationship to clinical progression.

The spatial transcriptomics data give a partial answer to whether the CCT2 relationship exists outside the dish. In postmortem frontal cortex, with spots enriched for layer 5b projection neuron markers (62), CCT2 expression across ALS samples tracks the cell cycle axis score closely, and the correlation is unchanged when CCT2 is removed from the axis itself. So the relationship between CCT2 and the cell cycle program is present in end-stage human cortex and is not an artifact of CCT2 being counted on both sides. At the group level, CCT2 is lower in ALS cortex than in control. That is consistent with what we see in the cells. About three quarters of sporadic patients, those in S-ALS-1 and S-ALS-2, have CCT2 below control in purified motor neurons, so part of the reduction in bulk tissue is real rather than compositional. Neuronal loss will also contribute, since CCT2 is highly expressed in neurons and the tissue has fewer of them. What bulk tissue cannot show is the S-ALS-3 subgroup moving the other way, which is the same averaging problem that makes the cohort-level comparison uninformative. This is the same reason the primary analysis was done in purified motor neurons in the first place. Two things limit this analysis. The number of motor cortex samples available was too small to use, so the result is from frontal cortex, where the enriched cells are cortical projection neurons rather than the spinal motor neurons the iPSC-MNs model, and spatial data cannot tell us which subtype a given donor would have belonged to during life. It is a test of one relationship, not a test of the subtypes.

Several limitations should be kept in mind. The first is culture composition. The AnswerALS differentiations vary in purity, and subtype assignment is related to differentiation quality, most strongly for S-ALS-1. We checked this by re-deriving the subtypes after adjusting for composition markers and by re-estimating the CCT2 association under those adjustments. The subtype labels move more under adjustment than the CCT2 expression difference does, which changed very little. S-ALS-1 should be read with more caution than the other two subtypes, and its overlap with ALS-Glia may be partly a composition effect on both sides. The association between CCT2 and ALSFRS-R decline, however, weakened under the same adjustment and was no longer significant. The second is replication. There is no independent iPSC motor neuron cohort of comparable size in which we could test the three-axis structure, and the comparisons to Target ALS bulk tissue and to GSE56500 are directional only. The Bye et al. cohort, generated at a different site with a different differentiation protocol and with survival data, is the natural test of whether CCT2 identifies a subset of patients with faster disease, and it is under controlled access at the time of writing (47). CCT2 also shows why any replication will need to be stratified. It differs sharply between subtypes within this cohort and barely at all between sporadic patients and controls, because the subtypes move in opposite directions and average out. The third is proteomics depth, which is enough to confirm the CCT2 direction but not to build a subtype signature at the protein level. Finally, all of the mechanism in this paper is inferred from expression. We have not manipulated MYC, mTORC1 or CCT2 in these cells, and the claims about what the S-ALS-3 program does to a motor neuron are hypotheses rather than results.

## Conclusion

Sporadic ALS motor neurons from living patients can be grouped into three regulatory states along metabolic, cell cycle and inflammatory axes, and one of these states can be identified by CCT2 alone. The states are not genetic. The CCT2 marker holds at the protein level, relates weakly to clinical progression, and its relationship to the cell cycle program is visible in end-stage human cortex. The next steps are clear enough: replicate the structure in an independent iPSC cohort, test whether CCT2-high patients differ in survival, and find out in the cells what MYC and mTORC1 activity is doing to a motor neuron. If the subtypes hold up, they give a way to enroll patients in trials by biology rather than by diagnosis, which is what the field has been asking for.

## Software and reproducibility

Analyses were run in R. Every reported result derives from a pinned analysis object, and each analysis script reproduces its own landmark values on a fresh session before writing output.

## Supporting information

lasso

genomic

clinical

axiscomp

axisgenes

## Acknowledgement

The Los Angeles Claude D. Pepper Older Americans Independence Center (NIA P30AG094848).

## Notes

### Competing Interest Statement

The authors have declared no competing interest.

